# Lasting effects of repeated Δ9-tetrahydrocannabinol (THC) vapor inhalation during adolescence in male and female rats

**DOI:** 10.1101/426064

**Authors:** Jacques D. Nguyen, K. M. Creehan, Tony M. Kerr, Michael A. Taffe

## Abstract

Adolescents are regularly exposed to Δ^9^-tetrahydrocannabinol (THC) via smoking, and, more recently, vaping, cannabis / extracts. Growing legalization of cannabis for medical and recreational purposes, combined with decreasing perceptions of harm, makes it increasingly important to determine the consequences of frequent adolescent exposure for motivated behavior and lasting tolerance in response to THC. Male and female rats inhaled THC vapor, or that from the propylene glycol (PG) vehicle, twice daily for 30 minutes from postnatal day (PND) 35-39 and PND 42-45 using an e-cigarette system. Thermoregulatory responses to vapor inhalation were assessed by radio-telemetry during adolescence and from PND 86-94; chow intake was assessed in adulthood. Blood samples were obtained from additional adolescent groups following initial THC inhalation and after four days of twice daily exposure. Additional groups exposed repeatedly to THC or PG during adolescence were evaluated for intravenous self-administration of oxycodone as adults. Female, not male, adolescents developed tolerance to the hypothermic effects of THC inhalation in the first week of repeated exposure despite similar plasma THC levels. Each sex exhibited tolerance to THC hypothermia in adulthood after repeated adolescent THC with THC greater potency exhibited in females. Repeated-THC male rats consumed more food than their PG treated control group, in the absence of a significant bodyweight difference. Adolescent THC did not alter oxycodone self-administration in either sex, but increased fentanyl self-administration in females. Repeated THC vapor inhalation in adolescent rats results in lasting consequences observable in adulthood.

**Abbreviations:** PG, propylene glycol; THC, Δ^9^tetrahydrocannabinol;

## Introduction

Significant numbers of adolescents are exposed to Δ^9^-tetrahydrocannabinol (THC) on a regular basis via the smoking and, more recently, vaping, of cannabis and/or cannabis extracts. Epidemiological data confirm that 5-6% of 12^th^ grade students in the USA use cannabis nearly daily, and 13.9% have used in the past month (Miech et al., 2018). About 10% of 12^th^ grade students have *vaped* cannabis at least once in the past year and 5% in the past month (*ibid*). Furthermore, growing legalization of cannabis use for medical and recreational purposes, and a decreasing perception of harm, suggests these populations will only grow in coming years. Human epidemiological evidence poses clear limitations for interpretation and it is therefore critical to determine the consequences of frequent adolescent exposure to THC in well-controlled, and translationally valid, animal models.

Repeated injection of THC in adolescent rats produces lasting effects in adulthood, including decreased bodyweight (Rubino et al., 2008), impaired spatial working memory (Rubino et al., 2009), increased heroin self-administration (Ellgren et al., 2007), increased reinstatement of heroin seeking (Stopponi et al., 2014) and greater sensitivity to learning impairments produced by THC (Winsauer et al., 2011). Most prior investigations have employed the injected route of administration (Cha et al., 2007; Ellgren et al., 2007; Rubino et al., 2008; Winsauer et al., 2011) which may not match the human condition very well. For example, the duration of effect of THC on hypothermia in the rat lasts hours longer following i.p. administration compared to a vapor inhalation regimen that produces a similar temperature nadir and peak plasma THC levels (Nguyen et al., 2016; Taffe et al., 2015). The majority of human use of cannabis is via inhalation which entails a comparatively rapid onset and offset with a shorter overall duration of activity. Thus, the study of inhaled delivery of cannabis constituents in rodent models may improve translational inferences.

The <u>first goal</u> of this study was to determine if vapor inhalation of THC reduces body temperature in the adolescent rat, as this is a key measure of THC activity in rodents. Our e-cigarette based model has been validated previously in adult rats, producing THC-typical effects on nociception and body temperature (Javadi-Paydar et al., 2018; Nguyen et al., 2016) and plasma THC levels comparable to those reached by human marijuana users (Hartman et al., 2015; Huestis et al., 1992). We have not shown efficacy in adolescent rats which is a critical gap in the validation of this model and therefore an important goal. The <u>second, and major, goal</u> was to determine if twice daily THC exposure over consecutive days during adolescence produces tolerance, indicative of a degree of THC exposure sufficient to induce lasting changes in the central nervous system. Our recent study in adult rats showed tolerance to the hypothermic and antinociceptive responses to THC after twice (female) or thrice (male or female) daily inhalation (Nguyen, J. D. et al., 2018a). The <u>third goal</u> was to test the hypothesis that the development of tolerance differs across rat sex, since prior work has shown that female rats develop tolerance more rapidly and at a lower mg/kg adjusted dose following twice-daily parenteral injection of THC (Wakley et al., 2014). Our prior work shows that male and female adult rats achieve similar plasma THC levels after identical inhalation conditions (Javadi-Paydar et al., 2018) but that adult female rats become tolerant with less intensive THC exposure compared with males (Nguyen, J. D. et al., 2018a). Thus, it was hypothesized that female adolescent rats would be more sensitive than the males, developing tolerance after fewer sessions and to a greater extent. The <u>fourth goal</u> was to determine if there were lasting consequences of adolescent THC inhalation in the adult rat in terms of 1) tolerance to acute THC exposure; 2) alterations in weight gain and feeding behavior (Sofia and Barry, 1974) and 3) in the propensity to self-administer oxycodone (Nguyen et al., 2019) or fentanyl, as has previously been reported for heroin (Ellgren et al., 2007; Stopponi et al., 2014).

## Methods

### Subjects

Female (N=40) and male (N=48) Wistar rats (Charles River, Livermore, CA) were shipped on postnatal day (PND) 19 and entered the laboratory on PND 22. Groups of 16 male and 16 female rats were used in the radio-telemetry experiments, groups of 24 male and 16 female rats were used in the self-administration experiments and groups of 8 male and 8 female rats were used in the plasma THC concentration assessments. Rats were housed in humidity and temperature-controlled (23±2 °C) vivaria on 12:12 hour light:dark cycles and all studies were conducted in the rats’ scotophase. Animals had *ad libitum* access to food and water in their home cages. Gelatin nutritional support (DietGel® Recovery, ClearH2O, Westbrook, ME, USA) was provided during weekdays from PND 32-46 given the age of the animals, the radio-telemetry surgery and the unfamiliar experimental apparatus. All procedures were conducted under protocols approved by the Institutional Care and Use Committee of The Scripps Research Institute in a manner consistent with the principles of the NIH Guide for the Care and Use of Laboratory Animals.

### Radio-telemetry

Rats (N=16 per sex) were implanted with sterile radio-telemetry transmitters (Data Sciences International, St Paul, MN; TA11TA-F20) in the abdominal cavity as previously described (Taffe et al., 2015; Wright et al., 2012) on PND 25. Animals were recorded in a dark testing room separate from the vivarium in either the vapor inhalation chambers (males) or in separate clean home cages (females) in the same room. We have previously shown that this moderate difference in procedure has negligible effect on the body temperature response to cannabinoids (Javadi-Paydar et al, 2018). This difference in procedure resulted in a 30 minute recording time point for male rats but not for female rats. Radio-telemetry transmissions were collected via telemetry receiver plates (Data Sciences International, St Paul, MN; RPC-1 or RMC-1) placed under the cages as described in prior investigations (Aarde et al., 2013; Miller et al., 2013; Wright et al., 2012). The sexes were treated identically up until PND 86 and thereafter there were some slight differences; the order and details of studies are outlined in **Table 1**.

**Table 1.**
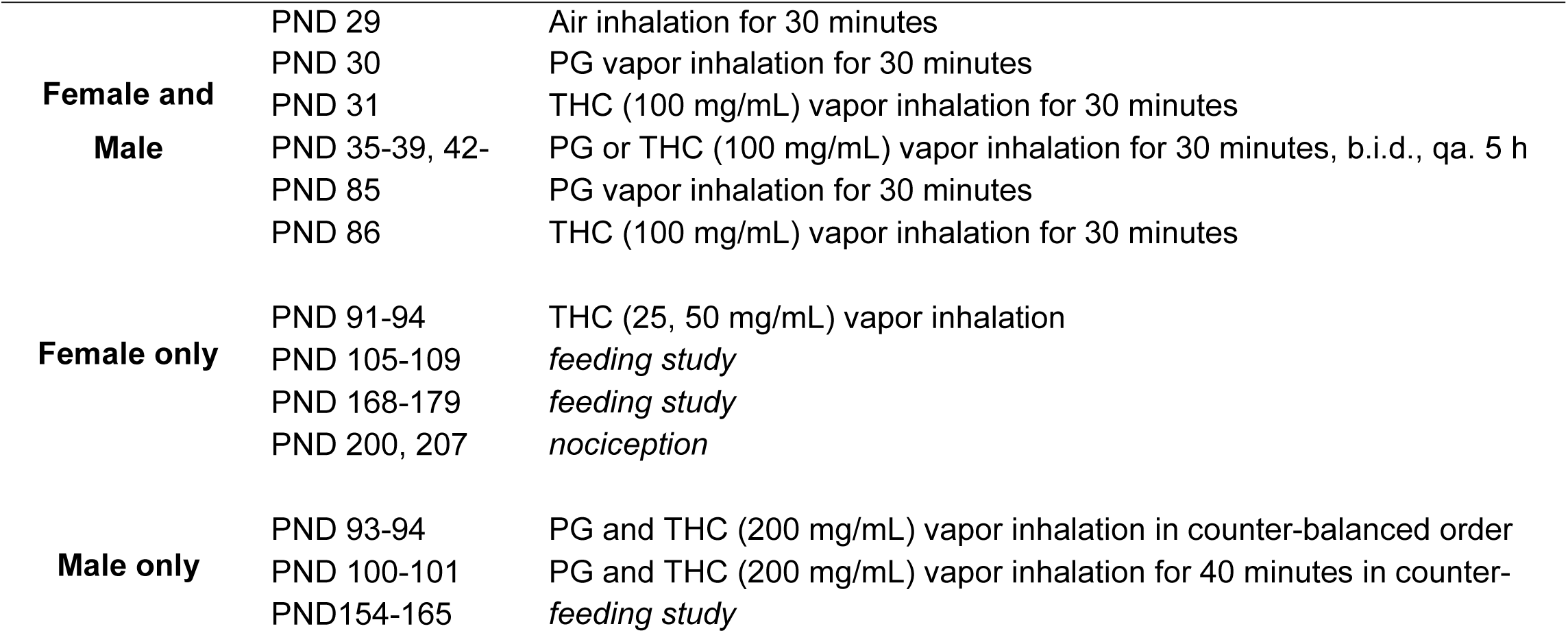
Order of inhalation and feeding studies for male and female radio-telemetry cohorts.

### Intravenous Catheterization

Rats (N=16 female; N= 24 male) were anesthetized with an isoflurane/oxygen vapor mixture (isoflurane 5% induction, 1-3% maintenance) and prepared with chronic indwelling intravenous catheters as described previously (Aarde et al., 2017; Miller et al., 2015; Nguyen et al., 2017) on PND 84-87, i.e. after adolescent vapor exposure (see below). Briefly, the intravenous catheters consisted of a 14.5-cm length of polyurethane based tubing (Micro-Renathane®, Braintree Scientific, Inc, Braintree, MA) fitted to a guide cannula (Plastics One, Roanoke, VA) curved at an angle and encased in dental cement anchored to an ∼3 cm circle of durable mesh. Catheter tubing was passed subcutaneously from the animal’s back to the right jugular vein. Catheter tubing was inserted into the vein and tied gently with suture thread. A liquid tissue adhesive was used to close the incisions (3M™ Vetbond™ Tissue Adhesive: 1469SB, 3M, St. Paul, MN). A minimum of 4 days was allowed for surgical recovery prior to starting an experiment. For the first three days of the recovery period, an antibiotic (cefazolin) and an analgesic (flunixin) were administered daily. During testing and training, intravenous catheters were flushed with ∼0.2-0.3 ml heparinized (166.7 USP/ml) saline before sessions and ∼0.2-0.3 ml heparinized saline containing cefazolin (100 mg/mL) after sessions. Catheter patency was assessed once a week after the last session of the week, via administration of the ultra-short-acting barbiturate anesthetic Brevital sodium (1% methohexital sodium; ∼0.2 ml (10 mg/ml); Eli Lilly, Indianapolis, IN) through the catheter. Animals with patent catheters exhibit prominent signs of anesthesia (pronounced loss of muscle tone) within 3 sec after infusion. Animals that failed to display these signs were considered to have faulty catheters, and if catheter patency failure was detected, data that were collected after the previous passing of this test were excluded from analysis.

### Drugs

Δ^9^-tetrahydrocannabinol (THC) was administered by vapor inhalation. Doses are varied, and therefore described, by altering the concentration in the propylene glycol (PG) vehicle, the puff schedule and the duration of inhalation sessions in this approach. The ethanolic THC stock was aliquoted in the appropriate volume, the ethanol evaporated off and the THC was then dissolved in the PG to achieve target concentrations. (-)-Oxycodone HCl and fentanyl citrate were dissolved in saline (0.9% NaCl), for injection. The THC was provided by the U.S. National Institute on Drug Abuse and the PG, oxycodone and fentanyl were obtained from Sigma-Aldrich Corporation (St. Louis, MO, USA).

### Inhalation procedure

The inhalation procedure followed methods that have been recently described (Javadi-Paydar et al., 2018; Nguyen et al., 2016). Sealed exposure chambers were modified from the 259mm × 234mm × 209mm Allentown, Inc (Allentown, NJ) rat cage to regulate airflow and the delivery of vaporized drug to rats. An e-vape controller (Model SSV-1; La Jolla Alcohol Research, Inc, La Jolla, CA, USA) was triggered to deliver the scheduled series of puffs from Protank 3 Atomizer (Kanger Tech; Shenzhen Kanger Technology Co.,LTD; Fuyong Town, Shenzhen, China) e-cigarette cartridges by a computerized controller designed by the equipment manufacturer (Control Cube 1; La Jolla Alcohol Research, Inc, La Jolla, CA, USA). Type 2 sealed exposure chambers (La Jolla Alcohol Research, Inc; La Jolla, CA, USA) and second generation e-vape controllers (Model SSV-2; La Jolla Alcohol Research, Inc, La Jolla, CA, USA) with Herakles Sub Ohm Tank e-cigarette cartridges (Sense; Shenzhen Sense Technology Co., LTD; Baoan Dist, Shenzhen, China) or Smok Baby Beast Brother TFV8 sub-ohm tank (with the V8 X-Baby M2 0.25 ohm coil; SMOKTech, Nanshan, Shenzhen, China), triggered by MedPC IV software (Med Associates, St. Albans, VT USA), were used for adolescent PG/THC exposure in the groups destined for the pharmacokinetic and self-administration experiments. The chamber air was vacuum-controlled by a chamber exhaust valve (i.e., a “pull” system) to flow room ambient air through an intake valve at ∼1 L per minute. This also functioned to ensure that vapor entered the chamber on each device triggering event. The vapor stream was integrated with the ambient air stream once triggered. For all studies the system delivered four 10-s vapor puffs, with 2-s intervals, every 5 minutes for 30 minutes (i.e., last puff at 25 minutes with 5 min inhalation time).

### Adolescent Vapor Exposure

Each experimental cohort (i.e., destined for radio-telemetry, intravenous self-administration or plasma collection experiments) was randomly divided into experimental (repeated adolescent THC inhalation) and control (repeated adolescent PG inhalation) subgroups, which received 30 minute episodes of vapor exposure, twice per day at a 5 h interval (∼2 and 7 h from the start of dark) to either THC (100 mg/mL) or the propylene glycol (PG) vehicle on sequential days PND 35-39 and again on PND 42-46 (plasma assessment groups received vapor exposure only for the first four days, see below). Plasma THC declines rapidly in the first hour after ceasing inhalation exposure and reaches low levels by 4 hours after vapor initiation (Javadi-Paydar et al., 2018; Nguyen et al., 2016). The experimental timeline for the radio-telemetry cohorts is outlined in Table 1. The self-administration groups received intravenous catheter implant surgery on PND 84-87 and initiated the intravenous self-administration (IVSA) of oxycodone (0.15 mg/kg/infusion; 8 h sessions; Fixed Ratio 1 response contingency) on PND 112. Following an acquisition interval of 17 sessions the rats completed six sessions of Fixed Ratio (8 h) and Progressive Ratio (3 h) dose substitution (0.006, 0.06, 0.15 mg/kg/infusion) in a counter-balanced order. Treatment of the plasma concentration assessment cohorts is described below.

### Feeding procedure

To begin a test day, all chow was removed at the start of the dark cycle in the vivarium. Rats were moved to a procedure room, weighed and placed individually in separate rat cages equipped with J-style hanging chow dispensers (Guinea Pig Feeder; Ancare, Bellmore, NY, USA) starting 2 h into the dark period. Chow was removed and weighed every 2 hours for 6 hours; rats were weighed again at the end of the test interval. The feeding study was conducted PND 105-109 and 168-179 in the female radio-telemetry groups and PND 154-165 in the male radio-telemetry groups.

### Plasma THC Levels

Separate groups of 8 male and 8 female rats were received on PND 22. These rats received a single inhalation session (THC 100 mg/mL; 30 minutes) on PND 31, after which a blood sample (∼500 uL) was obtained by acute venipuncture under inhalation anesthesia. The rats then received twice daily inhalation sessions from PND 36-39 with a second blood sample obtained after the first session on PND 39. This timing was designed to capture the first session of complete tolerance observed in the female rats in the telemetry study, i.e., Day 4. Blood samples were also obtained from these groups on PND 86 following a THC 100 mg/mL (30 minutes) inhalation session and on PND 100 and 107 following THC 50 or 200 mg/mL inhalation for 30 minutes in a counterbalanced order.

Blood samples were collected (∼500 ul) via jugular needle insertion under anesthesia with an isoflurane/oxygen vapor mixture (isoflurane 5% induction, 1–3% maintenance) 35 minutes post-initiation of vapor inhalation. Plasma THC content was quantified using fast liquid chromatography/mass spectrometry (LC/MS) adapted from (Irimia et al., 2015; Lacroix and Saussereau, 2012; Nguyen, J. D. et al., 2018a). 5 μL of plasma were mixed with 50 μL of deuterated internal standard (100 ng/mL cannabidiol (CBD)-d3 and THC-d3; Cerilliant), and cannabinoids were extracted into 300 μL acetonitrile and 600 μL of chloroform and then dried. Samples were reconstituted in 100 μL of an acetonitrile/methanol/water (2:1:1) mixture. Separation was performed on an Agilent LC1100 using an Eclipse XDB-C18 column (3.5um, 2.1mm × 100mm) using gradient elution with water and methanol, both with 0.2 % formic acid (300 μL/min; 73-90%). Cannabinoids were quantified using an Agilent MSD6140 single quadrupole using electrospray ionization and selected ion monitoring [CBD (m/z=315.2), CBD-d3 (m/z=318.3), THC (m/z=315.2) and THC-d3 (m/z=318.3)]. Although rats were not exposed to any CBD in this experiment, the assay included it for consistency and comparison of the plasma concentrations with other studies ongoing in the laboratory which include simultaneous analysis of CBD and THC. Calibration curves were conducted for each assay at a concentration range of 0-200 ng/mL and observed correlation coefficients were 0.999.

### Oxycodone Self-Administration

Intravenous self-administration was conducted in operant boxes (Med Associates) located inside sound-attenuating chambers, in an experimental room (ambient temperature 22 ± 1 °C; illuminated by red light) outside of the housing vivarium, starting on PND 112. To begin a session, the catheter fittings on the animals’ backs were connected to polyethylene tubing contained inside a protective spring suspended into the operant chamber from a liquid swivel attached to a balance arm. Each operant session started with the extension of two retractable levers into the chamber. Following each completion of the response requirement (response ratio), a white stimulus light (located above the reinforced lever) signaled delivery of the reinforcer and remained on during a 20-sec post-infusion timeout, during which responses were recorded but had no scheduled consequences. Drug infusions were delivered via syringe pump. The training dose (0.15 mg/kg/infusion; ∼0.1 ml/infusion) was selected from prior oxycodone self-administration studies (Nguyen et al, 2018; Wade et al, 2015). The rats were trained in 8 h sessions using a Fixed Ratio 1 (FR1) response contingency during weekdays (5 days per week) for 17 (male) or 16 (female) sessions. Thereafter the animals were assessed in a dose substitution procedure (with dose order counter-balanced within the groups) in 8 h sessions under a FR1 contingency. Following this, the dose substitution (with dose order counter-balanced within the groups) was completed under a Progressive Ratio (PR) contingency procedure. In the PR paradigm, the required response ratio was increased after each reinforcer delivery within a session (Hodos, 1961; Segal and Mandell, 1974) as determined by the following equation (rounded to the nearest integer): Response Ratio=5e^(injection number*j)–5 (Richardson and Roberts, 1996). The j value was set to 0.2 and sessions were terminated when 60 minutes elapsed without a response, or after 3 h. Female rats also completed two successive dose substitution tests under FR1 in 4 h sessions involving first fentanyl (0.0, 1.25, 2.5, 5.0 µg/kg/inf) and then fentanyl (0.0, 0.625, 2.5, 10.0 µg/kg/inf), each in a counterbalanced order, starting PND 200 (see **Supplemental Methods and Figure S6** for additional experiments completed prior to this experiment for these animals).

### Data Analysis

Body temperature and activity rate (counts per minute) were collected on a 5-minute schedule but are expressed as 30 min averages for analysis in the study. The time identifier is at the end of the interval, i.e., the 60 minute timepoint is the average from 35-60 minutes after the start of inhalation. The time courses for data collection are expressed relative to the start of the inhalation. Any missing temperature values were interpolated from the values before and after the lost time point, this typically involves fewer than 10% of observations. Missing activity rate datapoints were not replaced because values can change dramatically from one 5-min interval to another, thus there is no rational basis for interpolating. The 60 minute timepoint was selected for follow-up comparison across sexes and / or treatment conditions because our prior work (Javadi-Paydar et al., 2018; Nguyen et al., 2016; Nguyen, J. D. et al., 2018a) found that this time is consistently when the maximum hypothermia is observed after THC inhalation.

Statistical analysis of temperature, activity, plasma THC concentrations, bodyweight, infusions earned, and lever discrimination was conducted with Analysis of Variance (ANOVA). Between-groups factors included Sex or Adolescent treatment condition, as appropriate. Within-subjects factors of Session, Dose, Postnatal day (PND) and Vapor inhalation condition were included as appropriate. Significant main effects or effects of the interaction of factors were followed with post hoc analysis using Tukey (multi-level factors) or Sidak (two-level factors) correction for multiple comparisons. A p-value < 0.05 was used as the criterion for a significant result. These analytical approaches are consistent with recommendations on experimental design and analysis in pharmacology (Curtis et al., 2015). All analysis used Prism 6, 7 or 8 for Windows (v. 6.02, 7.03, 8.1.1; GraphPad Software, Inc, San Diego CA).

## Results

### THC-induced hypothermia in adolescent rats

The initial experiment confirmed that vapor inhalation of THC induces hypothermia in adolescent rats of each sex (**Figure 1A, B**). Furthermore, the male and female groups destined for repeated-THC versus repeated-PG were not different in response to PG on PND 30 and to THC on PND 31. The three-factor analysis confirmed significant effects of Vapor inhalation condition [F(1,252)=310.1; *p*<0.05], of Time after vapor initiation [F(8,252)=12.58; *p*<0.05] and the interaction of Time with Vapor inhalation condition [F(8, 252)=16.13; *p*<0.05] in female rats; there were no significant main effects or interactions with the Group of eventual assignment. The three-way analysis for the male groups confirmed significant effects of Vapor inhalation condition [F(1,280)=215.1; *p*<0.05], of Time after vapor initiation [F(9,280)=16.19; *p*<0.05] and the interaction of Time with Vapor inhalation condition [F(9,280)=10.61; *p*<0.05] and the interaction of the Group of eventual assignment with Vapor inhalation condition [F(1,280)=10.22; *p*<0.05]. Post hoc analysis of this latter interaction failed to confirm any differences between groups destined for repeated-PG and repeated-THC in either PG or THC inhalation conditions.

**Figure 1:**
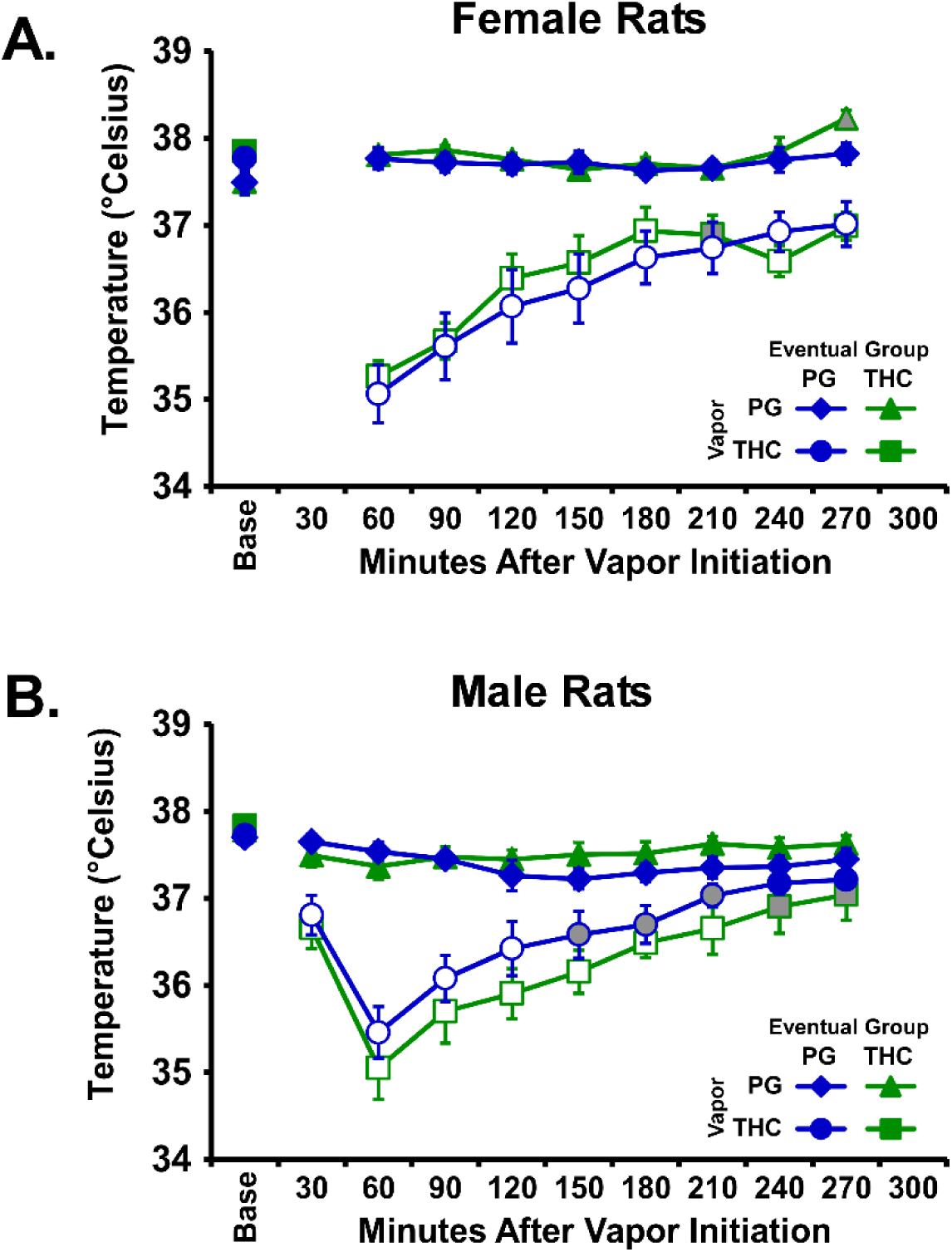
Mean (N=8 per group; ±SEM) body temperature responses after inhalation of PG (on PND30) or THC (100mg/mL; on PND31) vapor for 30 minutes in the subgroups eventually assigned to the repeated-PG or repeated-THC A) female and B) male groups. Open symbols indicate a significant difference from both vehicle at a given time-point and the within-treatment baseline (Base), while shaded symbols indicate a significant difference from the baseline only.

Follow up analysis of body temperature 60 minutes after the start of inhalation including the repeated-PG and repeated-THC cohorts of each sex confirmed a main effect of vapor inhalation condition [F(1,28)=249.1; *p*<0.05] but not of group or of the interaction. The post hoc test confirmed a significantly lower temperature after THC vapor compared with PG within each group and did not confirm any differences across all the groups within PG or THC vapor inhalation conditions.

### Effect of repeated THC or PG inhalation on hypothermia in adolescent rats

There were no changes in the body temperature response of the repeated-PG groups across the recording interval for any of the recording days (**Supplemental Figure S1**), but both sexes in the repeated-THC groups became hypothermic following inhalation during each of the chronic exposure weeks (**Figure 2**). Tolerance to the hypothermic effects differed between the sexes. Statistical analysis of the female animals’ temperature confirmed significant effects of Time after vapor initiation and Day in Week 1 [Time: F(9,63)=32.61; *p*<0.05; Day: F(5,35)=18.72; *p*<0.05; Interaction: F(45,315)=11.39; *p*<0.05] and Week 2 [Time: F(9,63)=9.69; *p*<0.05; Day: n.s.; Interaction: F(45,315)=3.12; *p*<0.05]. Statistical analysis of the male animals’ temperature confirmed significant effects in Week 1 [Time: F(10,70)=31.26; *p*<0.05; Day: F(5,35)=4.35; *p*<0.05; Interaction: F(50,350)=4.72; *p*<0.05] and Week 2 [Time: F(10,70)=31.62; *p*<0.05; Day: F(5,35)=6.467; *p*<0.05; Interaction: F(50,350)=4.28; *p*<0.05]. The post hoc test confirmed that a progressive tolerance to hypothermia developed in the female rats across the first 4 days of exposure with no hypothermia observed on days 4 and 5. The post hoc test also confirmed that significant tolerance developed in Days 3-5 relative to Day 1 in the males, although this was limited to the 120-180 minute time points. In Week 2, the post hoc test confirmed a significant reduction in body temperature relative to the PG day 60 minutes after the start of inhalation on Days 7-9 for the female rats and Days 6-10 for the male rats. Follow up analysis was conducted on the temperature recorded 60 minutes after the start of inhalation for repeated-THC cohorts of each sex to directly compare the magnitude of hypothermia across days and sex (**Figure 3**). The ANOVA confirmed a main effect of day [F(11,154)=19.14; *p*<0.05] of Sex [F(1,14)=10.6; *p*<0.05] and of the interaction [F(11,154)=6.36; *p*<0.05]. The post hoc test further confirmed that significant reductions in temperature following THC inhalation (relative to the PG condition) were observed for each THC session in male rats.

**Figure 2:**
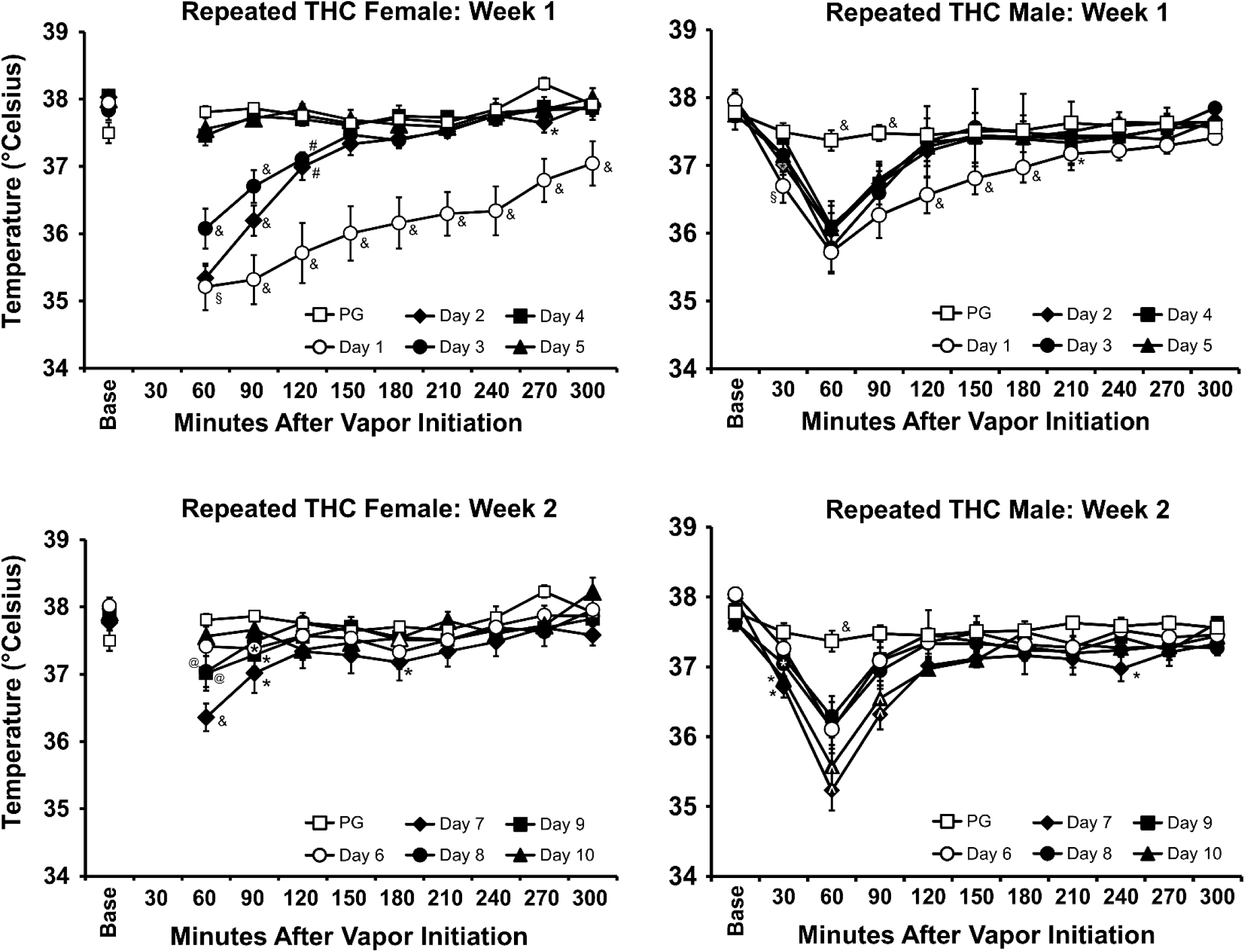
Mean (N=8 per sex; ±SEM) body temperature recorded for the first vapor inhalation session of each day of the repeated treatment weeks is depicted for the repeated THC groups; Week 1 = PND 35-39; Week 2 = PND 42-46. The PND 30 PG inhalation data are depicted in upper and lower panels for comparison. The pre-inhalation baseline temperature is indicated by “Base”. A significant difference from all other days at a given time after the start of inhalation is depicted by &, a significant difference from all days except Day 2 by §, a significant difference from PG, Day 4 and 5 by #, a significant difference from PG, Day 7 and 10 by @, a significant difference from PG, Days 6, 8 and 9 by ^, and a significant difference from PG by *.

**Figure 3:**
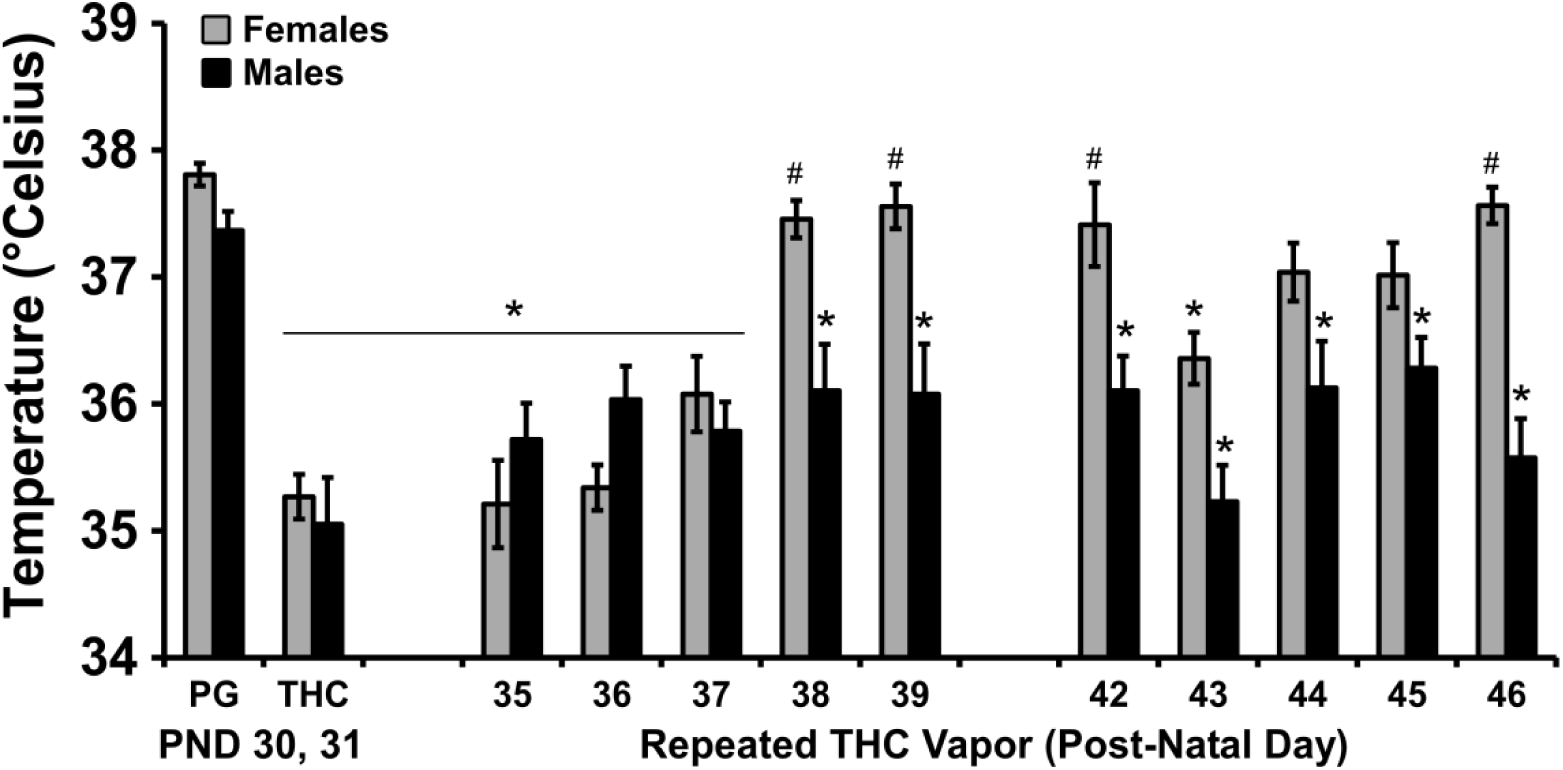
Mean (N=8 per sex; ±SEM) body temperature 60 minutes after the start of inhalation for all inhalation days during adolescence. A significant difference from the PG value, within group, is indicated with * and a significant sex difference, on a given day, by #.

Significant reductions in body temperature were also confirmed for female rats on PND days 31, 35-37, and 43; a significant sex difference was confirmed for PND 38-42 and 46. A reduction in body temperature was observed during the first daily session of all 10 days of repeated THC inhalation but no change in body temperature was observed during PG inhalation. Body weight was most significantly reduced by repeated THC during the adolescent weeks in male rats (see **Supplemental Figure S3**).

### Adult THC inhalation

Tolerance to the hypothermic effects of THC inhalation were observed in adulthood in the repeated-THC groups as compared with their respective repeated-PG control groups (**Figure 4**). The statistical analysis of the temperature of the repeated PG females after inhalation during adulthood confirmed a significant effect of Time after vapor initiation [F(8,56)=13.63; *p*<0.05], of Vapor Concentration condition [F(3,21)=20.87; *p*<0.05] and of the interaction of Time with Vapor Concentration condition [F(24,168)=2.85; *p*<0.05]. Analysis of the temperature of the repeated THC females confirmed a significant effect of Time after vapor initiation [F(8,56)=8.05; *p*<0.05], of Vapor Concentration condition [F(3,21)=13.93; *p*<0.05] and of the interaction of factors[F(24,168)=6.29; *p*<0.05]. The analysis of the temperature of the repeated PG males confirmed a significant effect of Time after vapor initiation [F(9,63)=18.24; *p*<0.05] and of the interaction of Time with Vapor Concentration condition [F(18,126)=6.75; *p*<0.05]. Analysis of the temperature of the repeated THC males confirmed a significant effect of Time after vapor initiation [F(9,63)=11.35; *p*<0.05], of Vapor Concentration condition [F(2,14)=4.34; *p*<0.05] and of the interaction of factors [F(18,126)=5.08; *p*<0.05].

**Figure 4:**
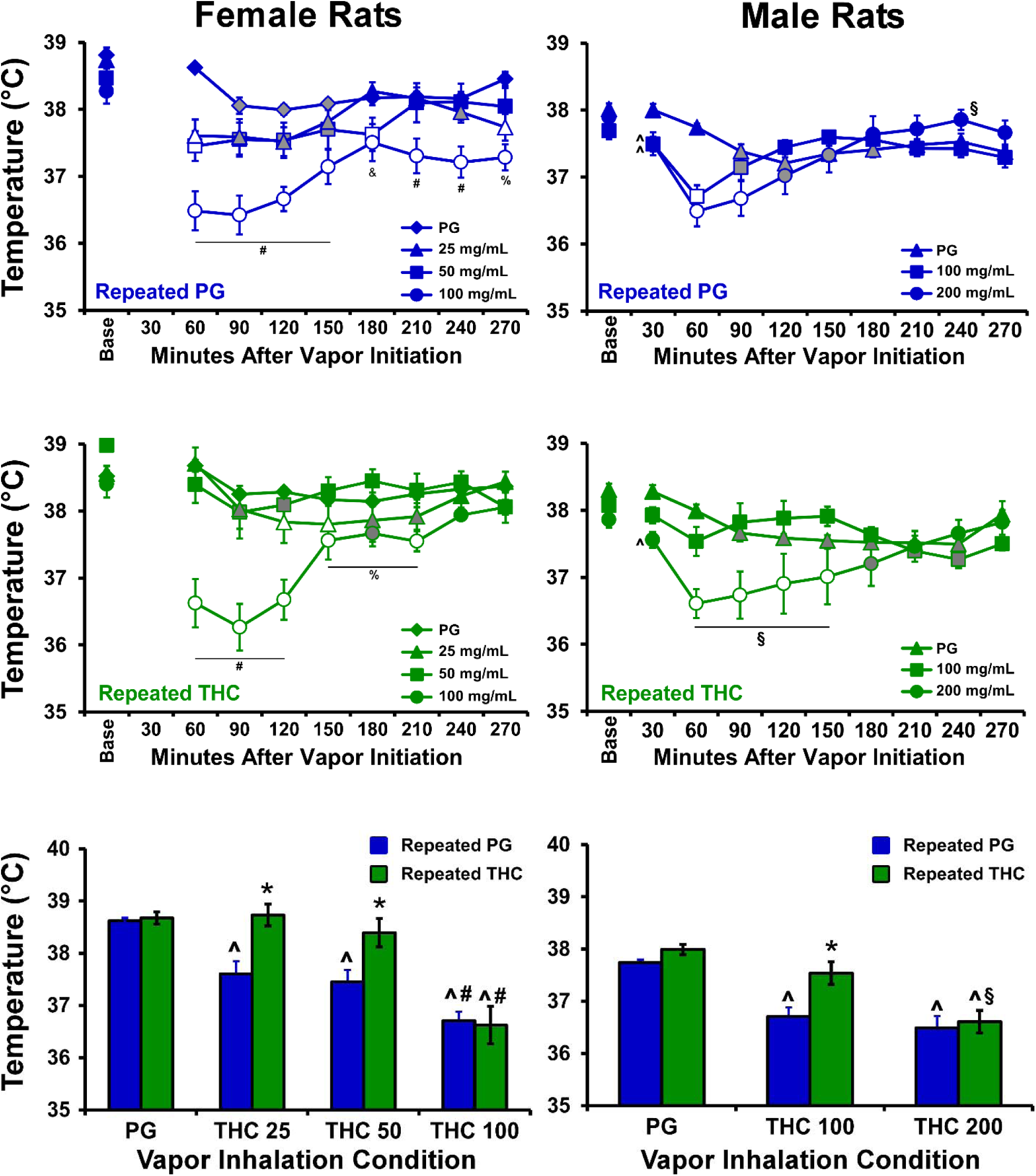
Mean (±SEM) body temperature of female and male rat cohorts (N=8 per group) exposed to repeated PG or THC during adolescence and challenged in acute sessions with vapor from PG or varying concentrations of THC (25-200 mg/mL) from PND 85-94. Open symbols indicate a significant difference from the pre-inhalation baseline (Base) and the corresponding time after PG inhalation, and grey symbols represent a difference from the baseline only. A significant difference from the PG condition is indicated with ^, from the 25 and 50 mg/mL condition with #, from the 100 mg/mL condition by §, from the 50 mg/mL condition by % and from the 25 mg/mL condition by &. A significant difference between treatment groups at given dose, within sex, is indicated with *.

Analysis of the temperature for the 60 minute interval in the female groups confirmed a significant effect of Group [F(1,14)=5.57; *p*<0.05], of Vapor Inhalation condition [F(3,42)=41.96; *p*<0.05] and of the interaction of factors [F(3,42)=3.89; *p*<0.05] on body temperature. The post hoc analysis confirmed a group difference following inhalation of THC 25-50 mg/mL. The analysis of temperature for the 60 min interval for the male groups likewise confirmed a significant effect of Group [F(1,14)=6.44; *p*<0.05] and of Vapor Inhalation condition [F(2,28)=31.27; *p*<0.05], but not of the interaction of factors. The post hoc test confirmed a significant difference between the groups in the THC 100 mg/mL condition.

### Plasma THC Levels

The plasma THC concentrations did not differ between samples obtained from adolescent male and female rats in either the acute (PND 31) or chronic (PND 39) THC inhalation experiments (**Figure 5**). The analysis confirmed a significant effect of Acute versus Chronic experiment phase [F(1,14)=26.64; *p*<0.05] without significant effect of Sex or of the interaction of factors. During adulthood, plasma THC levels varied with Vapor Inhalation condition [F(2,26)=20.94; *p*<0.05] and Sex [F(1,13)=7.05; *p*<0.05]. The post hoc test confirmed a significant sex difference at the 50 mg/mL THC concentration and a difference from the 50 mg/mL concentration after 100 or 200 mg/mL inhalation for each sex.

**Figure 5:**
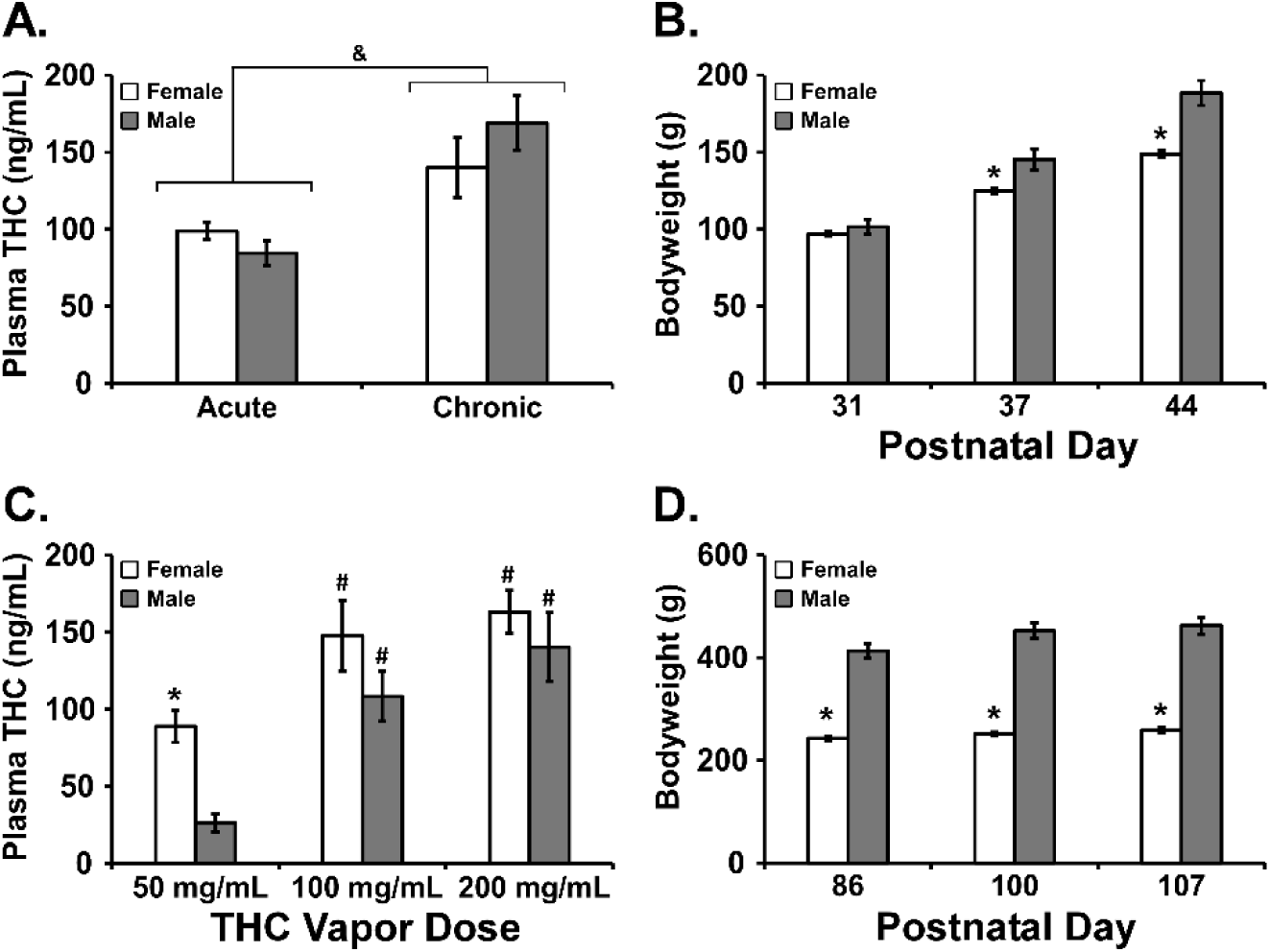
A, C) Mean (±SEM) plasma THC levels from female and male rats (N=8 per group) exposed to A) THC (100 mg/mL) vapor on PND 31 (Acute), 36-39 (Chronic) or C) THC (50-200 mg/mL) on PND 86, 100, 107. Corresponding body weights are presented for the adolescent (B) and adult (D) age intervals. A significant sex difference for a given dose or day is indicated with *, a difference from the Acute condition across groups is indicated with & and a difference from the 50 mg/mL condition with #.

Analysis of the bodyweights confirmed significant differences in the adolescent [PND: F(2,28)=1126; *p*<0.05; Sex: F(1,14)=10.19; *p*<0.05; Interaction: F(2,28)=71.21; *p*<0.05] and adult [PND: F(2,28)=183.8; *p*<0.05; Sex: F(1,14)=143.5; *p*<0.05; Interaction: F(2,28)=55.82; *p*<0.05] age ranges. The post hoc tests confirmed that there were significant sex differences in weight across all three days of the adolescent and adult age ranges.

### Adult food consumption

Feeding was assessed in 6 h sessions during PND 154-158 and PND 161-165 in the male rats and during PND 105-109, PND 168-172 and PND 175-179 in the female rats (**Figure 6**). One of the female rats from the original THC group was in apparently unrelated ill health and was not included in the PND 168-179 feeding study. No significant difference in food intake was confirmed for the female rats in any of the feeding studies although, interestingly, the female repeated-THC group remained slightly less sensitive to antinociceptive effects of THC when assessed from PND 200-207 (**Supplemental Figure S4**). The male THC group consumed more chow than the male PG Group with the analysis confirming significant effects of Group [F(1,14)=17.16; *p*<0.05], of Day [F(9,126)=4.78; *p*<0.05] and of the interaction of Group with Day [F(9,126)=4.97; *p*<0.05]. The post hoc test further confirmed the repeated-THC group consumed more food on PND 154, 155, 161, 162 and 165. Body weight prior to the session was not significantly different between the Groups for either sex.

**Figure 6:**
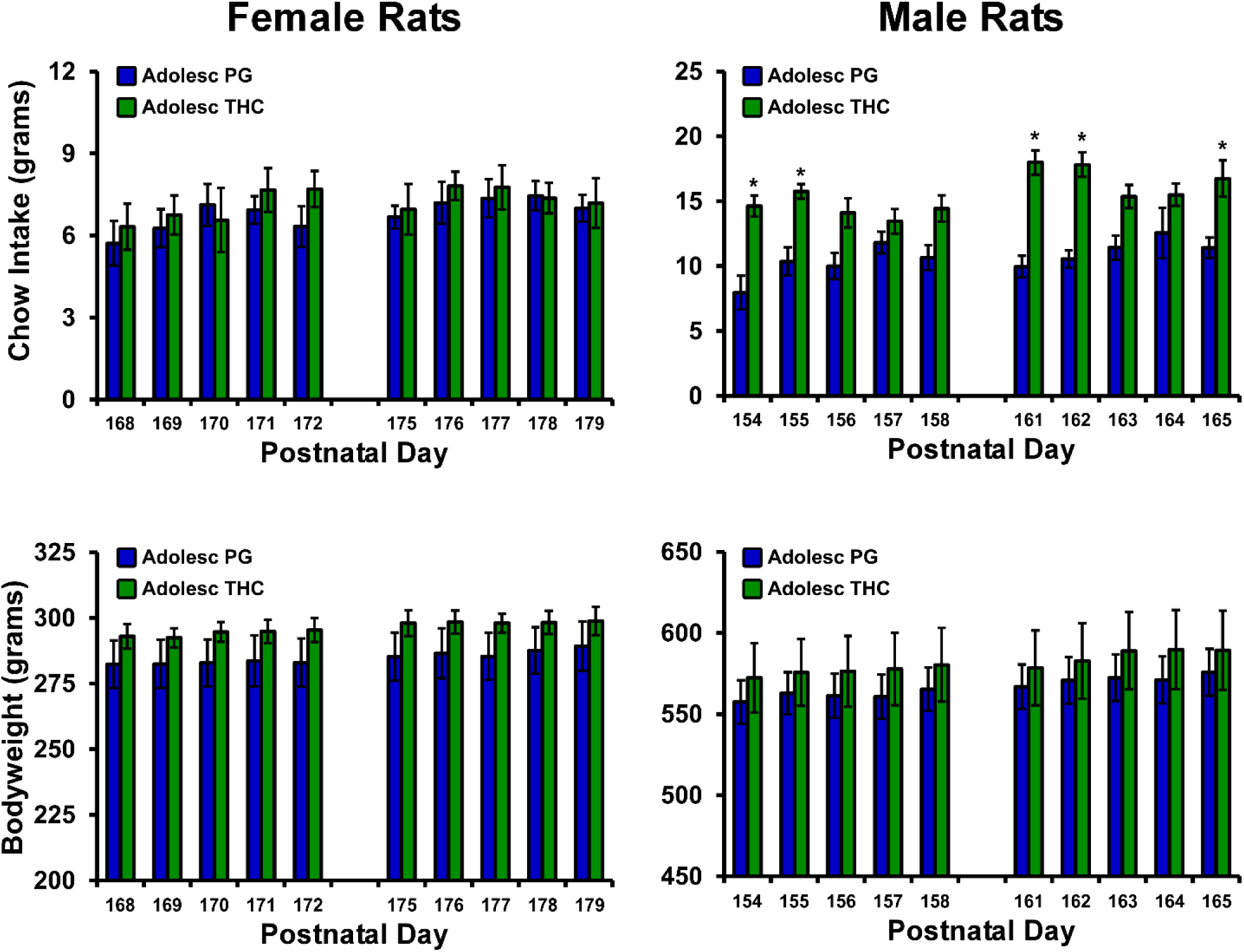
Mean (±SEM) chow intake and body weight of female and male rat cohorts exposed to repeated PG or THC during adolescence (N=8 per group except N=7 female/THC) and assessed during adulthood. A significant difference between treatment groups, within sex, is indicated with *.

### Oxycodone self-administration in adulthood

The male rats that were exposed to repeated vapor inhalation of either THC or PG vehicle during adolescence significantly escalated their oxycodone self-administration during 17 sessions of acquisition training as adults [F(16,352)=20.37; *p*<0.05]; there was no significant effect of Group (**Figure 7A**). Both groups exhibited a significant increase in appropriate drug-lever responding (lever discrimination) across sessions (**Figure 7B**) as the ANOVA confirmed significant effects of Session [F(16,352)=13.21; *p*<0.05] and of the interaction Group x Session interaction [F(16, 352)=2.053; *p*<0.05]. Analysis of oxycodone infusions during dose substitution experiments under a FR1 schedule confirmed a significant effect of Dose [F(2, 42)=12.46; *p*<0.05] but no effect of Group (**Figure 7C**). Similarly, analysis of the breakpoints under PR confirmed a significant effect of Dose [F(2, 42)=11.31; *p*=0.05] but no effect of Group (**Figure 7D**).

**Figure 7.**
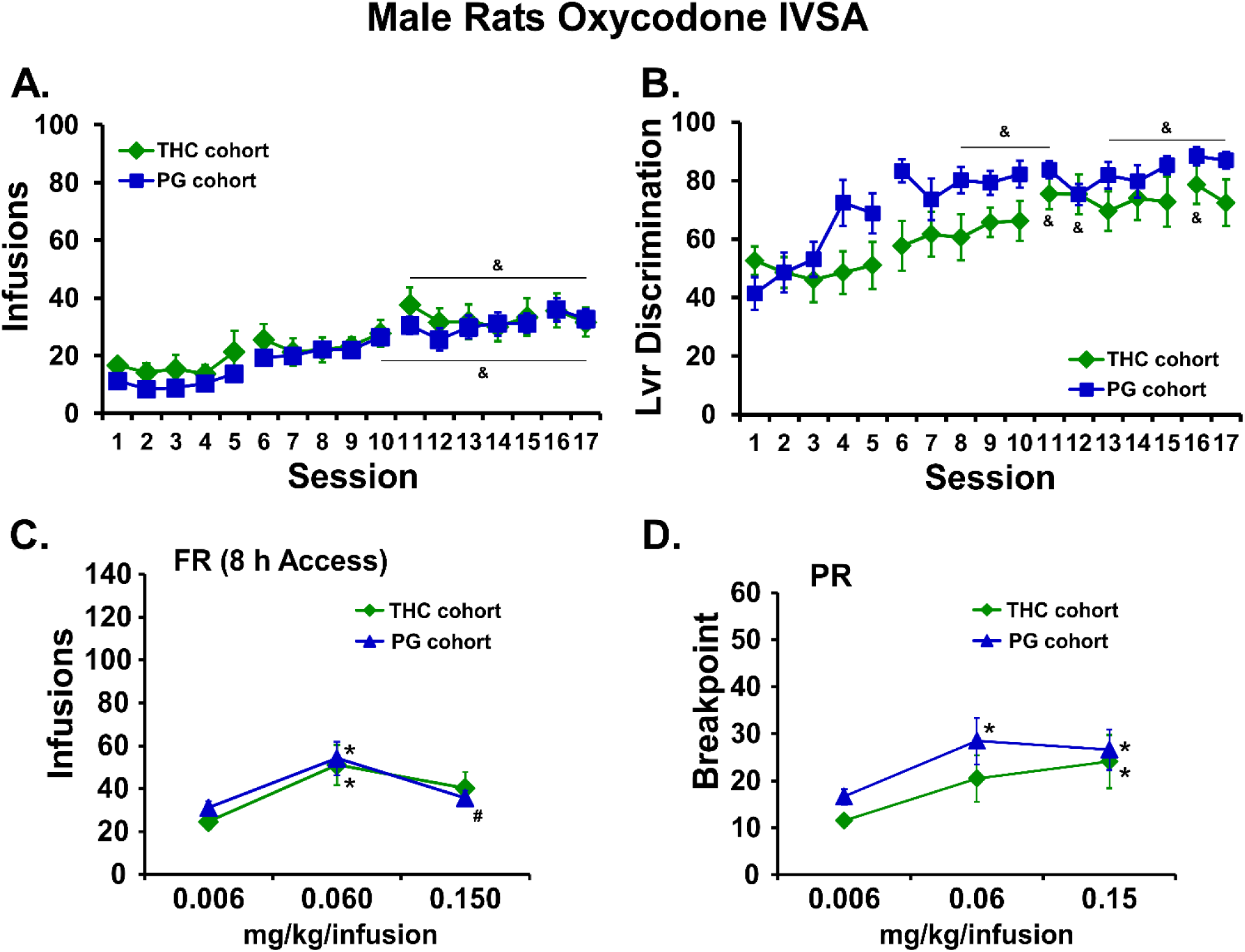
A) Mean infusions and B) percent drug-appropriate lever responding (Lvr Discrimination) for male rats trained to self-administer oxycodone (0.15 mg/kg/inf) within 8 h extended access sessions, starting on PND112. C) Mean (THC cohort, N=11; PG cohort, N=12; ±SEM) infusions during self-administration under a FR1 schedule and D) breakpoint values during self-administration under a PR schedule following acute injection of THC (0.006-0.15 mg/kg, i.p.). A significant difference from the first session is indicated by &. A significant difference from 0.006 mg/kg/infusion is indicated with * and a difference from the 0.06 is indicated with #.

The female rats that were exposed to repeated vapor inhalation of either THC or PG vehicle during adolescence also significantly escalated their oxycodone self-administration during the initial 16 sessions of acquisition training as adults [F(15,210)=9.96; *p*<0.05]; there was no significant effect of Group (**Figure 8A**). Both groups exhibited a significant increase in appropriate drug-lever responding (lever discrimination) across sessions (**Figure 8B**) as the ANOVA confirmed significant effects of Session [F(15,210)=5.09; *p*<0.05]. Analysis of oxycodone infusions during dose substitution experiments under a FR1 schedule confirmed a significant effect of Dose [F(2,26)=29.54; *p*<0.05] but no effect of Group (**Figure 8C**). Similarly, analysis of the breakpoints under PR confirmed a significant effect of Dose [F(2,26)=8.71; *p*<0.05] but no effect of Group (**Figure 8D**).

**Figure 8.**
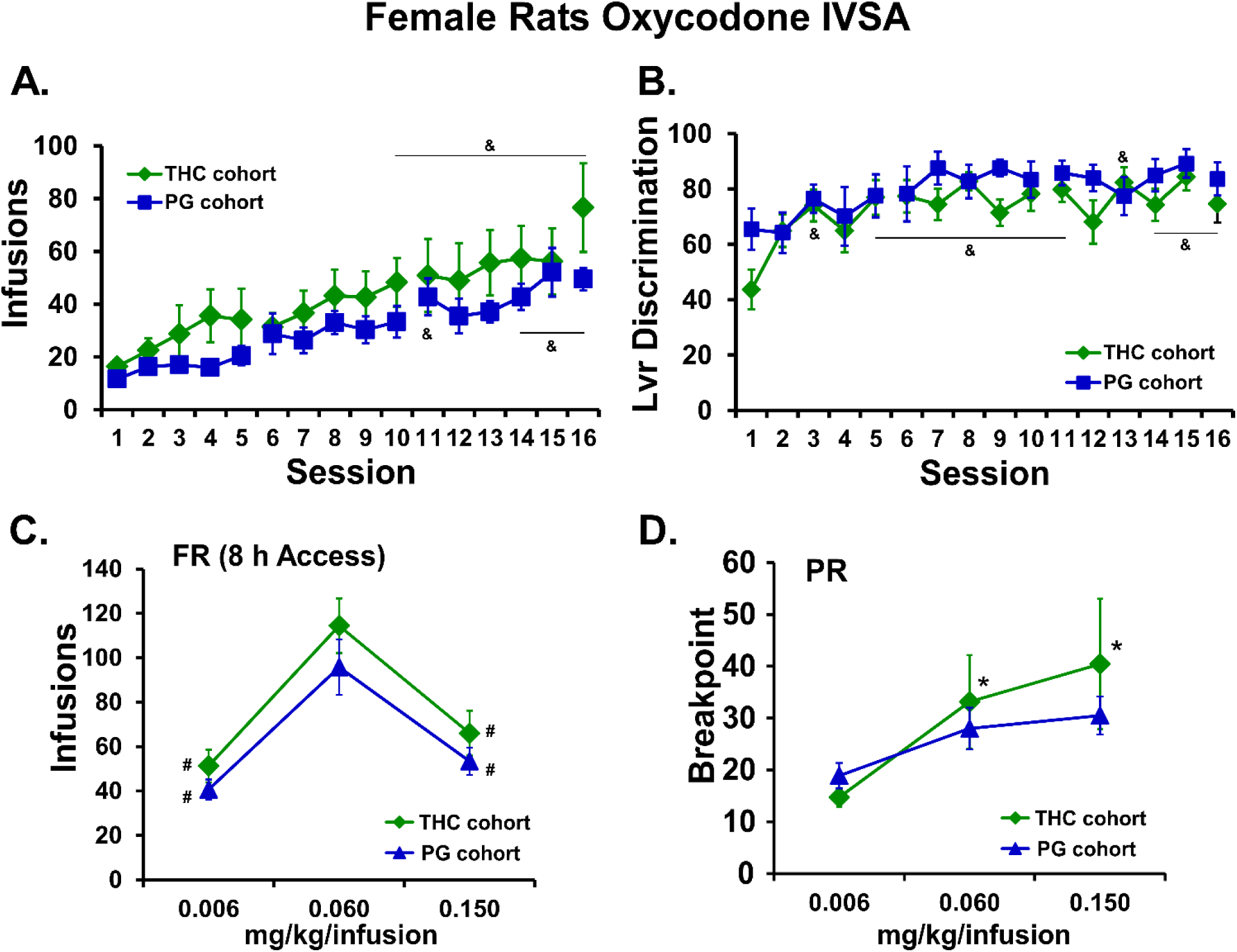
A) Mean (N=8; ±SEM) infusions and B) percent drug-appropriate lever responding (Lvr Discrimination) for female rats trained to self-administer oxycodone (0.15 mg/kg/inf) within 8 h extended access sessions, starting on PND112. C) Mean (THC cohort, N=7; PG cohort, N=8; ±SEM) infusions during self-administration under a FR1 schedule and D) breakpoint values during self-administration under a PR schedule following acute injection of THC (0.006-0.15 mg/kg, i.p.). A significant difference from the first session is indicated by &. A significant difference from 0.006 mg/kg/infusion is indicated with * and a difference from the 0.06 is indicated with #.

### Fentanyl self-administration in female rats

Preliminary analysis of the two fentanyl dose substitution experiments in the female rats (the catheters of N=6 repeated-THC remained patent) found no difference in the two overlapping doses in the first and second series. Therefore, the average of the vehicle and 2.5 ug/kg/infusion doses were used for formal analysis of the entire dose range. The analysis confirmed that the THC exposed female rats self-administered more fentanyl (significant effect of adolescent Treatment: F(1,72)=6.54; *p*<0.05; and of Dose: F(5,72)=17.38; *p*<0.0001) and the post hoc test confirmed a group difference at the 0.625 µg/kg/inf dose (**Figure 9**). Re-calculation of the oxycodone FR dose substitution excluding the THC-exposed animal who was not patent for the fentanyl study resulted in no change of the statistical significances.

**Figure 9.**
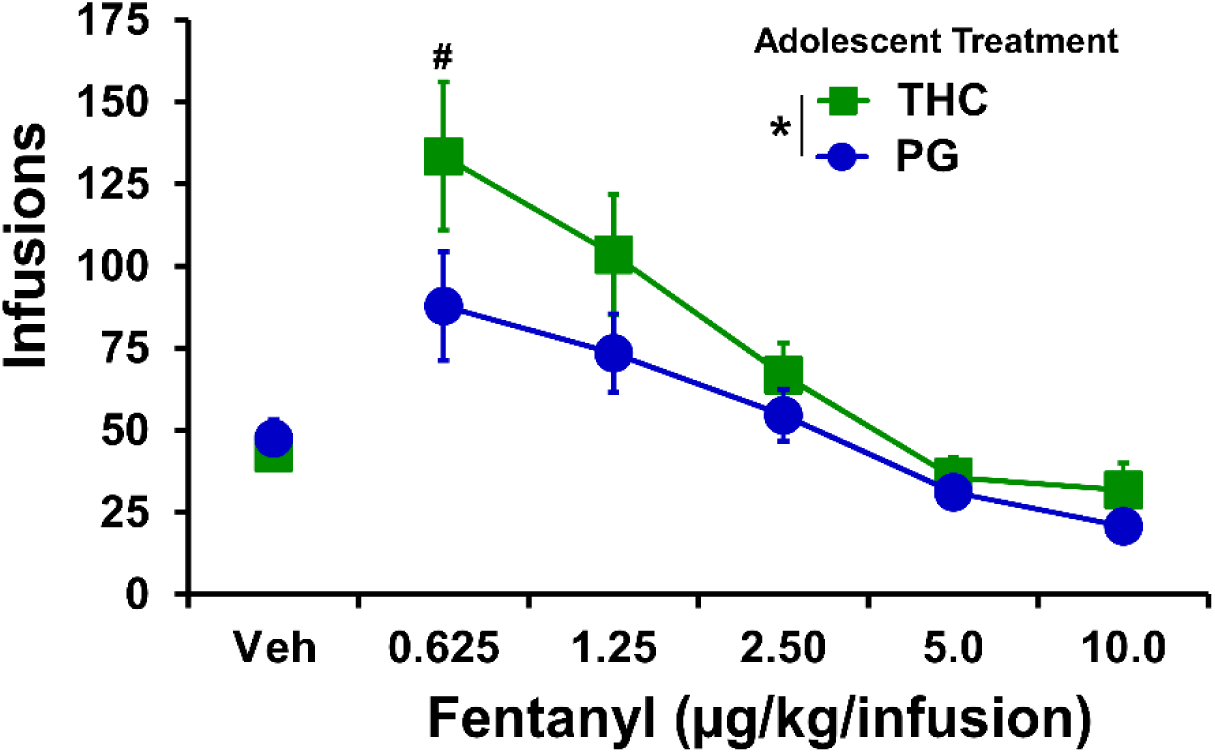
Mean (adolescent repeated-THC cohort, N=6; adolescent repeated-PG cohort, N=8; ±SEM) infusions of fentanyl obtained by female rats during self-administration under a FR1 schedule. A significant main effect of group (across doses) is indicated by * and a significant\ group difference at a specific dose with #.

### Sex differences in oxycodone self-administration

Follow-up analysis of the apparent sex difference in oxycodone self-administration, collapsed across adolescent treatment, confirmed that the female rats obtained more infusions during acquisition than did the males [Sex: F(1,37)=8.87; *p*<0.01; Session: F(15,555)=28.51; *p*<0.0001]; Interaction: F(15,555)=2.42; *p*=0.0021] and the post hoc test confirmed a significant difference between the sexes on sessions 14-16 (**Figure 10A**). Lever discrimination was also affected by sex [Sex: F(1,37)=2.67; *p*=0.1106; Session: F(15,555)=12.99; *p*<0.0001; Interaction: F(15,555)=1.93; *p*<0.05], however the post hoc confirmed a sex difference only in Session 3 (**Figure 10B**). This secondary analysis also confirmed that female rats self-administered more oxycodone in the FR procedure (**Figure 10C**) compared with male rats [Sex: F(1,36)=27.40; *p*<0.0001; Dose: F(2,72)=47.15; *p*<0.0001; Interaction: F(2,72)=8.44; *p*<0.001]. The post hoc test confirmed a sex difference at the 0.06 and 0.15 mg/kg/infusion doses. The post hoc test of the marginal mean for Dose confirmed significant differences in the infusions obtained in all three dose conditions. In the PR dose substitution (**Figure 10D**), there were no significant effects of sex [Sex: F(1,36)=2.10; *p*=0.1556; Dose: F(2,72)=19.11; *p*<0.0001; Interaction: F(2,72)=0.94; *p*=0.3939] confirmed. The post hoc test of the marginal mean for Dose confirmed significant differences in the infusions obtained in the 0.06 and 0.15 mg/kg/infusion condition compared with the 0.006 mg/kg/infusion dose condition. See **Supplemental Figure S5** for depiction of the self-administration data for all adolescent treatment groups, i.e., collapsed across sex.

**Figure 10:**
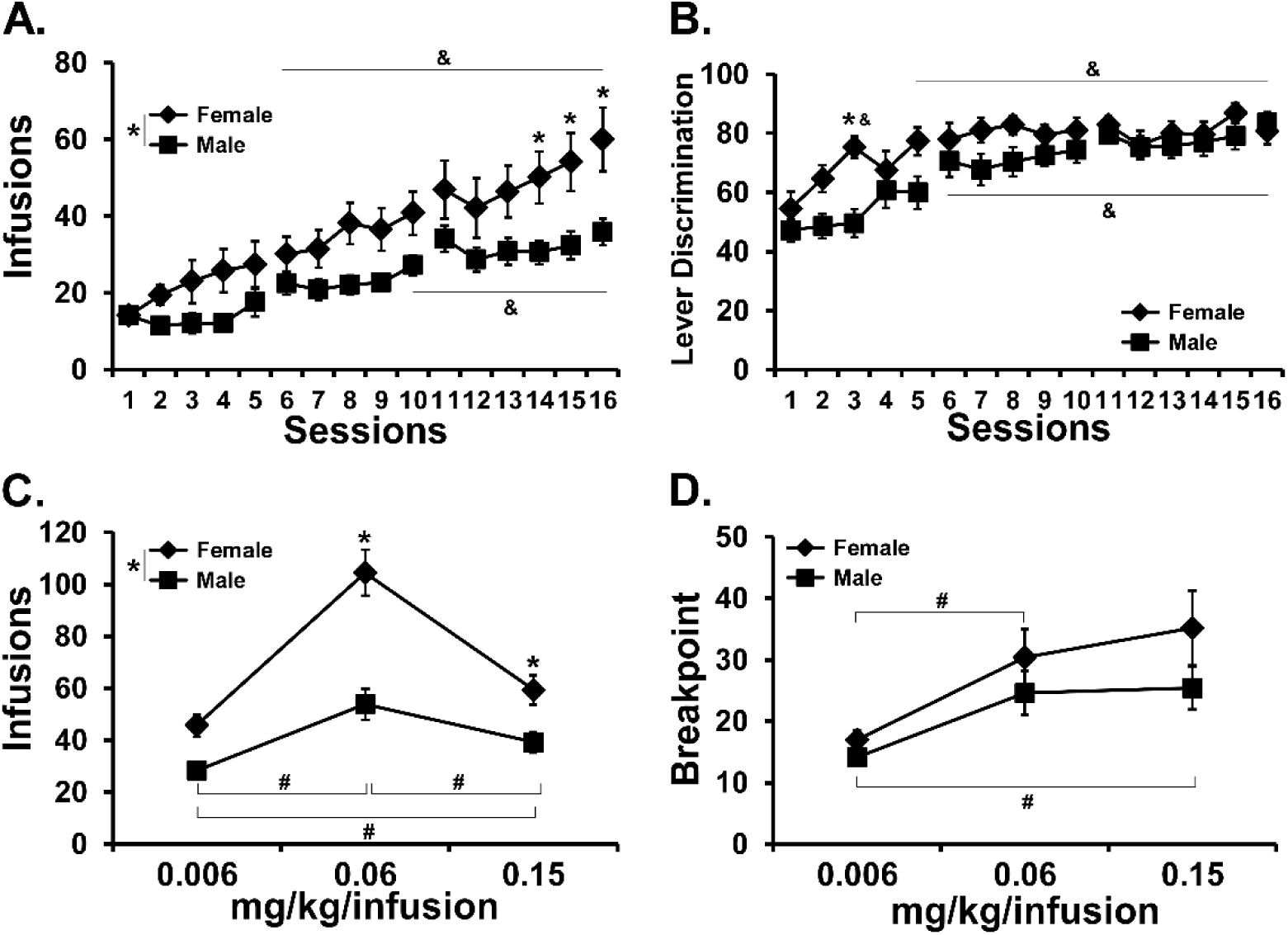
A) Mean (±SEM) infusions and B) lever discrimination during acquisition for the female (N= 15) and male (N= 24) animals collapsed across adolescent-treatment groups. C) Mean (±SEM) infusions obtained during the FR dose substitution and D) breakpoints reached in the PR dose substitution procedures are also depicted. A significant difference between sexes is indicated with *, a significant change from the first session within group by & and a significant difference between doses, collapsed across sex, is indicated with #.

## Discussion

This study showed that an e-cigarette-based method of Δ^9^-tetrahydrocannabinol (THC) inhalation produces hypothermia in adolescent rats, and tolerance with repeated exposure. Adolescent rats of each sex became hypothermic after vapor inhalation of THC, however, repeated inhalation produced rapid tolerance only in the females and a persisting tolerance in both sexes when assessed as adults. This was observed after two weeks (M-F) of twice daily exposure to an inhalation regimen which produced similar plasma levels of THC across the sexes (**Figure 5A**), thus confirming the enhanced sensitivity of female rats to the acute development of tolerance given a similar exposure to THC. In addition, tolerance to the thermoregulatory effects of inhaled THC lasted long past the chronic regimen, since adults of each sex from the repeated-THC groups were less sensitive to THC-induced hypothermia compared with their respective repeated-PG control groups when evaluated on PND 86 (**Figure 4**). The tolerance was dose-specific in each sex since an increase in the THC concentration resulted in a similar hypothermia in each adolescent treatment Group. Relatedly, there was a sex difference in the tolerance threshold during adulthood, i.e., the concentration of THC which produced minimal hypothermia in the repeated-THC group but a significant response in the repeated-PG group. This dose-effect difference may have been due to a sex difference in plasma THC concentrations produced during adulthood, particularly at lower effective dosing conditions (**Figure 5C**).

The THC-induced temperature change in the repeated-PG groups was lesser on PND 86 compared with PND 31, potentially due to the maturation of the rats and improved thermoregulation. This interpretation is consistent with the fact that body temperature observed 60 min after the start of inhalation on PND 86 was nearly identical to that observed in groups of naïve adult Wistar rats of the respective sexes following 30 min inhalation of THC (100 mg/mL) with no exposure during adolescence (Javadi-Paydar et al., 2018). As with our prior studies (Javadi-Paydar et al., 2018; Nguyen et al., 2016), locomotor behavior was not consistently affected by repeated THC. THC vapor inhalation increased activity rates slightly for about 30 min after inhalation ceased, and some tolerance to this appeared to be present in the female rats during the second week (**Figure S2**).

These results are qualitatively congruent with prior findings using parenteral injections or inhalation of THC. For example, adult female rats developed tolerance to THC to a greater extent than males, even with a lower per-injection dose (Wakley et al., 2015). Tolerance to the acute locomotor stimulant effects of injected THC developed more rapidly in adolescent female rats compared with adult male rats (Wiley et al., 2011) and locomotor tolerance to THC injections was approximately equivalent in adolescent male and female rats after 9 days of repeated THC injection (Wiley and Burston, 2014). Our prior work found female adult rats became tolerant after a repeated THC vapor inhalation regimen that did not produce tolerance in male adults (Nguyen, J. D. et al., 2018a). Adult male rats (females were not assessed) exposed to repeated marijuana smoke did not exhibit cross-tolerance to the locomotor suppressing effects of the endogenous cannabinoid agonist anandamide (Bruijnzeel et al., 2016). The finding of significantly lower bodyweight in the males during the second treatment week (**Figure S3**) is similar to evidence that repeated THC injections during adolescence reduced weight gain in male and female Wistar and Long-Evans rats in a 14 day treatment interval (Keeley et al., 2015) and in male Wistar rats during an 8 day repeated-injection study (Sofia and Barry, 1974). As a minor caveat, this study held age of drug exposure constant across sex, given a desire for face validity and the fact that human onset of marijuana use across sexes is better matched to chronological age rather than pubertal onset (Crane et al., 2015; Miech et al., 2018). Nevertheless, it would be of interest to compare a wider set of age ranges in future studies to further determine the role of pubertal events in any sex-mediated differences.

The present findings are likely mechanistically attributable, in part, to plasticity in the expression and/or function of the endogenous cannabinoid receptor subtype 1 (CB_1_). One prior study of repeated adolescent THC exposure found decreased CB_1_ receptor expression in the hippocampus of female, but not male rats (Weed et al., 2016). Another study found greater CB_1_ receptor desensitization in adolescents, compared with adults, and in female adolescents compared with male adolescents, following repeated THC injection. Changes were found in several brain regions including, importantly for the present thermoregulatory data, hypothalamus (Burston et al., 2010). One reason that adolescent female rats may be more sensitive to developing tolerance at a similar dose, brain or plasma THC level is that an active THC metabolite, 11-OH-THC, reaches higher concentrations in brain, compared with male rats (Wiley and Burston, 2014).

There were no apparent effects of adolescent THC exposure on the intravenous self-administration (IVSA) of oxycodone in either male or female rats. Analyzed across sex (**Figure S5**) there were no treatment-group differences in FR or PR dose substitution or infusions obtained during acquisition, however the THC exposed animals increased their lever-discrimination more slowly. There was a significant sex difference in acquisition and the FR dose-effect (**Figure 10**) which contrasts with one prior report (Mavrikaki et al., 2017), however this did not interact with adolescent treatment. The Mavrikaki et al. (2017) study used one-hour access sessions, thus the extended access model used here might be the reason for the sex difference in the current study. Alternately, it has previously been reported that female rats self-administer more morphine, heroin or fentanyl than do male rats (Cicero et al., 2003; Klein et al., 1997), thus it is unclear if sex differences in oxycodone self-administration exist apart from the procedural differences across a very limited number of comparisons. The acquisition in the males was consistent with that observed using similar procedures in a group of experimentally naïve adult males (Nguyen et al., 2019). Unfortunately the majority of rat oxycodone IVSA studies published so far have been in male rats (Austin Zamarripa et al., 2018; Blackwood et al., 2019; Bossert et al., 2018; Jordan et al., 2019; Leri and Burns, 2005; Mavrikaki et al., 2019; Nawarawong et al., 2018; Neelakantan et al., 2017; Nguyen et al., 2019; Nguyen, J. D. et al., 2018b; Pravetoni et al., 2014; Townsend et al., 2017; Wade et al., 2015; You et al., 2018; You et al., 2017); one exception was a study in pregnant rats (Vassoler et al., 2018). Interestingly, when female animals were evaluated on a fentanyl dose substitution under an FR procedure, the THC exposed rats self-administered more drug at the lowest dose (**Figure 9**). Additional investigation of specific opioids, particularly those that differ in potency, might be warranted in future studies.

Lasting effects of adolescent THC exposure were sometimes, but not always, found in other studies. Prior studies identified decreased bodyweight (Rubino et al., 2008), impaired spatial working memory (Rubino et al., 2009), and greater sensitivity to THC on a learning task (Winsauer et al., 2011) after repeated adolescent THC exposure. In contrast, repeated THC injection (3.2 mg/kg; 8 days) during adolescence did not affect THC place or taste conditioning in adulthood (Wakeford et al., 2016). Some prior studies have reported lasting motivational consequences, however the two available studies regarding the effect of repeated adolescent THC exposure on heroin IVSA reached different conclusions. One study in male Wistar rats used a regimen of twice daily injection in an escalating sequence of 2.5 mg/kg from PND 35–37, 5 mg/kg from PND 38–41 and 10 mg/kg from PND 42–45 (Stopponi et al., 2014). Another study injected male Long-Evans rats with THC (1.5 mg/kg, i.p.) every third day from PND 28-49 (Ellgren et al., 2007). Ellgren and colleagues reported increased heroin IVSA during acquisition as well as in a post-acquisition dose substitution procedure under a FR1 contingency in the repeated-THC group when evaluated as adults. In contrast, Stopponi and colleagues found no differences in the acquisition of heroin IVSA. The present study is consistent with the latter study and future work could determine if intermittency of exposure, overall dose, specific opioid drug or some other factors are responsible for the differences. Due to limited prior study, and varying THC exposure regimens used, it is not possible to draw strong conclusions about the breadth or selectivity of lasting motivational consequences of repeated adolescent exposure at this time. Related to this, feeding behavior was altered by repeated THC since the males consumed more food in 6 h focal feeding sessions compared with the PG control group. It will be interesting to determine in future studies if this reflects a motivational change akin to that reported for heroin IVSA by Ellgren and colleagues.

## Conclusions

The e-cigarette inhalation method is effective for the repeated daily exposure of adolescent rats, generating physiologically significant THC exposure and plasma levels consistent with those reported for humans after marijuana smoking or vaping (Hartman et al., 2015; Huestis et al., 1992). The exposure produced tolerance in the female adolescents but not in the males, consistent with a sex difference found previously with repeated THC injections. There are also lasting consequences of adolescent exposure on the adult animal including thermoregulatory tolerance in each sex, nociceptive tolerance in female rats (**Figure S4**), a feeding phenotype in male rats and increased fentanyl IVSA in female rats. This new approach has many advantages including that it avoids the stress of repeated injection, entails improved face validity and may provide improved pharmacokinetic fidelity with human exposure patterns.

## Author Contributions

MAT and JDN designed the studies, with refinements contributed by KMC and TMK. JDN, KMC and TMK performed the research and conducted initial data analysis. JDN and MAT conducted statistical analysis of data, created figures and wrote the paper. All authors approved of the submitted version of the manuscript.

## Acknowledgements

This work was supported by USPHS grants R01 DA035482 (Taffe, PI) and R44 DA041967 (Cole, PI). The National Institutes of Health / NIDA had no direct influence on the design, conduct, analysis or decision to publication of the findings. LJARI likewise did not influence the study designs, the data analysis or the decision to publish findings. The authors are grateful to Shawn M. Aarde, Ph.D. for significant contributions to the invention and initial validation of the vapor inhalation method, to Mr. Howard Britton for prototyping inhalation equipment and to Eric Zorrilla, Ph.D., Allison Kreisler, Ph.D. and Sophia A. Vandewater for assistance with the feeding assay. This is manuscript #29759 from The Scripps Research Institute.

## Competing Interest

The authors declare no financial interests that would create a conflict of interest for these studies.

## Notes

#### Summary of Updates

This version has been updated with results from a new group of female rats. The new study examined oxycodone and fentanyl self-administration after repeated adolescent vapor exposure to THC.

