## Supplementary Materials for "Lasting effects of repeated Δ9-tetrahydrocannabinol (THC) vapor inhalation during adolescence in male and female rats"

Running Head: Repeated adolescent THC inhalation

### Supplemental Methods:

#### Nociception Assay:

Tail withdrawal antinociception was assessed using a water bath (Bransonic® CPXH Ultrasonic Baths, Danbury, CT) maintained at 52 °C. The latency to withdraw the tail was measured using a stopwatch and a cutoff of 15 seconds was used (Wakley and Craft, 2011; Wakley et al, 2014). Tail withdrawal was assessed starting 35 minutes after the initiation of inhalation of PG or THC (100 mg/mL) for 30 minutes with the conditions assessed in a balanced order one week apart on PND 200 and PND 207.

#### Female Group Treatment Between Oxycodone and Fentanyl Dose Substitution Experiments:

The female rats in the oxycodone self-administration groups were returned to IVSA of oxycodone (0.15 mg/kg/inf) under FR1 in 4 h sessions for 7 sessions after the PR oxycodone dose substitution reported in the main body of the paper. Next, animals completed sessions self-administered oxycodone (0.15 mg/kg/inf) under the PR response contingency. For these studies the rats were injected i.p. with THC (0.0, 1.0, 5.0, 10.0 mg/kg) in a counter-balanced order. These pre-treatment conditions were then repeated, with animals self-administering oxycodone (0.06 mg/kg/inf). Finally the rats completed a dose substitution study under PR with fentanyl (0.0, 1.25, 2.5 and 5.0 μg/kg/infusion).

### Supplemental Results:

#### Effect of repeated PG inhalation on hypothermia in adolescent rats:

There were no changes in the body temperature of the repeated-PG groups across the recording interval for any of the recording days (**Figure S1**).

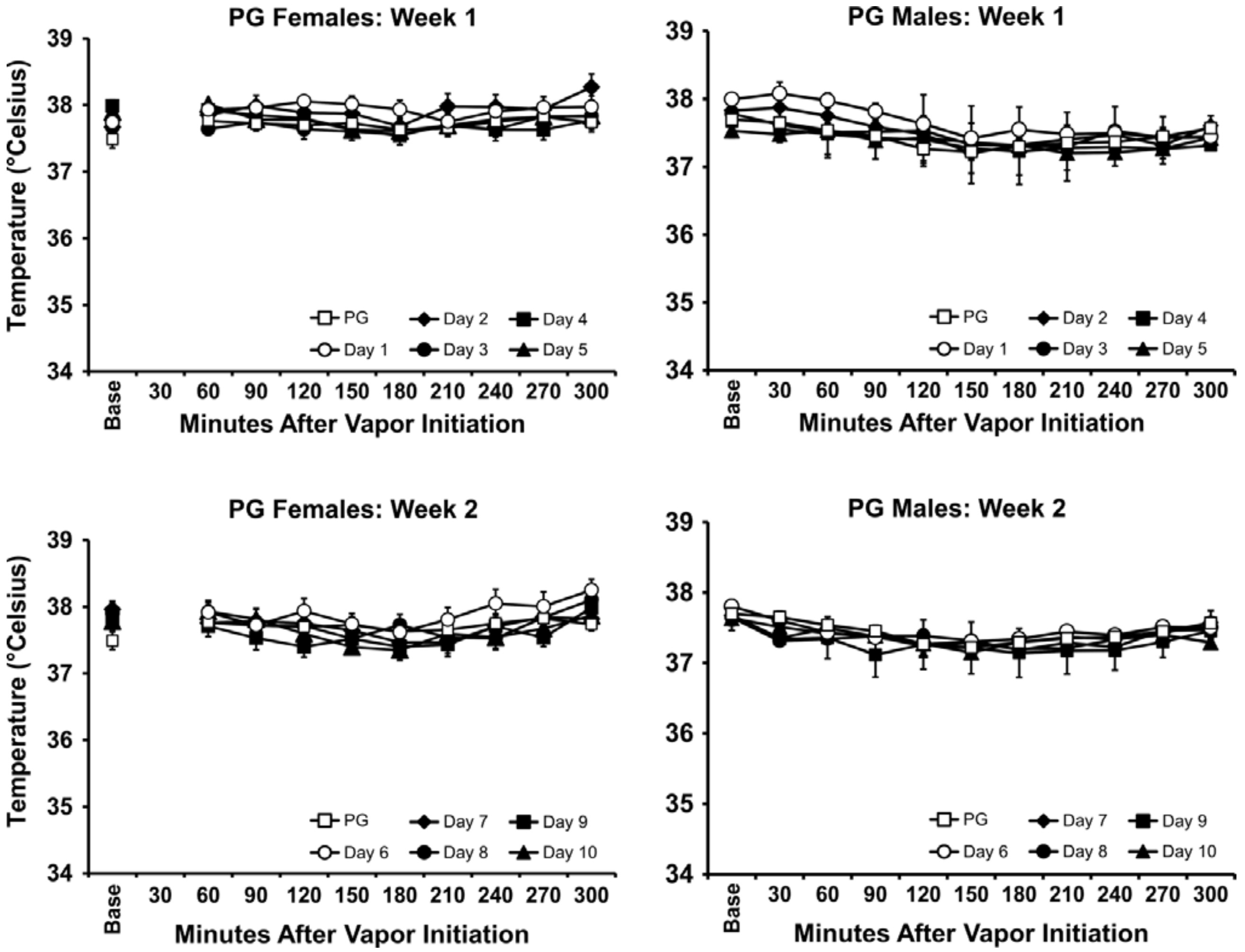


***Figure S1:*** *Mean (N=8 per sex; ±SEM) body temperature recorded for the first vapor inhalation session of each day is depicted for the repeated PG groups. The PND 30 PG data are depicted in upper and lower panels.*

**Effect of repeated THC inhalation on activity in adolescent rats:**

The analysis of the activity rates of the female rats exposed to repeated THC confirmed that both Time after vapor initiation and Day contributed to activity in Week 1 (Time: F(9,63)=16.87; *p*<0.0001; Day: F(5,35)=10.87; *p*<0.0001; Interaction: F(45,315)=5.62; *p*<0.0001) and Week 2 (Time: F(9,63)=22.18; *p*<0.0001; Day: F(5,35)=12.17; *p*<0.0001; Interaction: F(45,315)=5.39; *p*<0.0001). Unavoidable disruption in the room elevated activity levels 240 minutes after the start of inhalation on the PG day. Analysis of the male rats exposed to THC likewise confirmed significant effects on activity in Week 1 (Time: F(10,70)=33.72; *p*<0.0001; Interaction: F(50,350)=2.15; *p*<0.0001) and Week 2 (Time: F(10,70)=41.58;*p*<0.0001; Day: F(5,35)=7.48; *p*<0.0001; Interaction: F(50,350)=2.64; *p*<0.0001). Procedural differences (females were transferred from the recording cages into the vapor chambers for the inhalation interval and then returned)

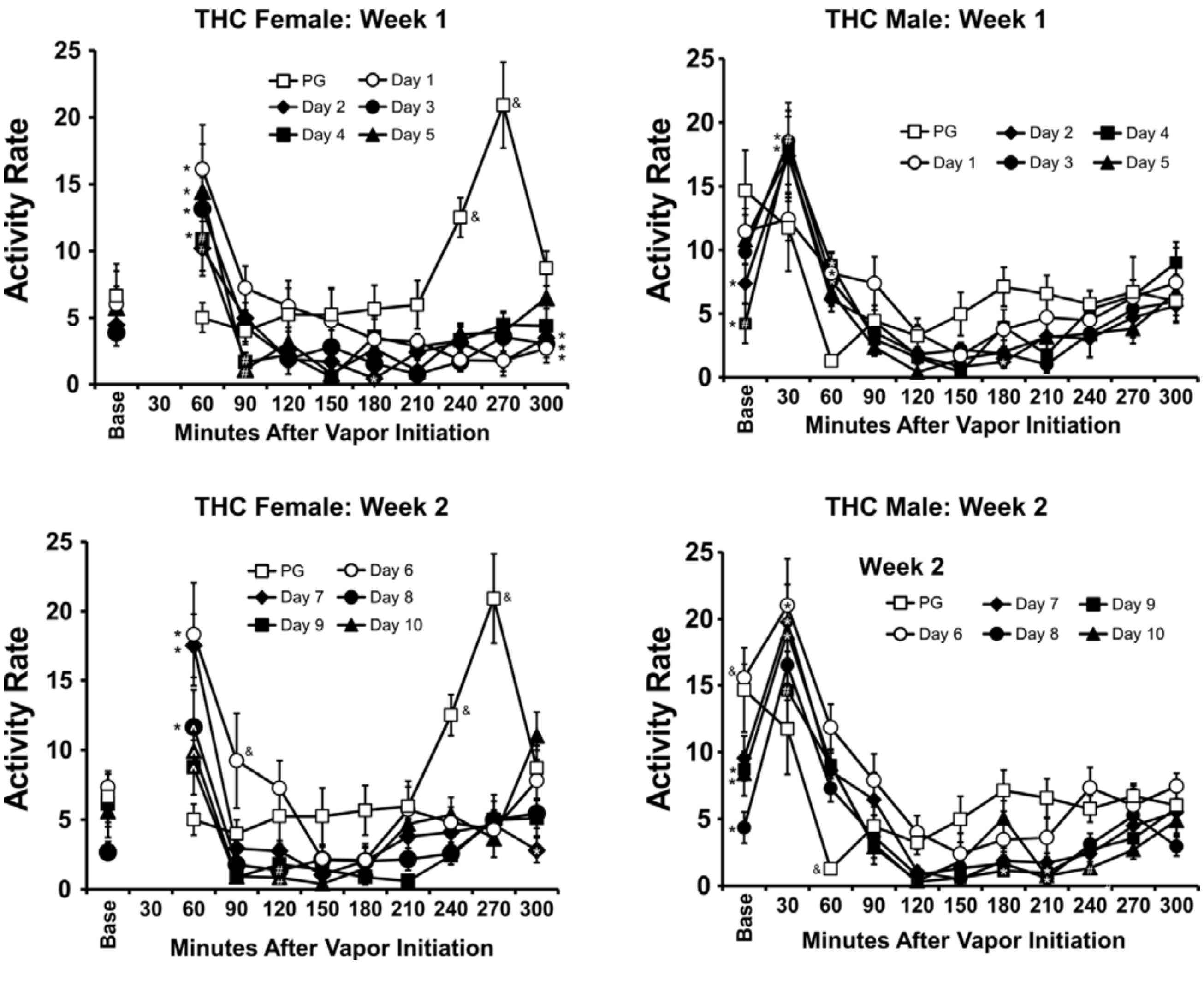


***Figure S2:*** *Mean (N=8 per sex; ±SEM) activity rate (counts/minute) recorded for the first vapor inhalation session of each day is depicted for the repeated THC groups. The PND30 PG data are depicted in upper and lower panels. A significant difference from all other days at a given time after the start of inhalation is depicted by &, a significant difference from PG and the first day of the week by #, a significant difference from Days 6 and 7 by ^, and a significant difference from PG by *.*

make it difficult to compare activity rates across sexes. If anything, THC vapor inhalation appeared to increase activity slightly for about 30 minutes after inhalation ceased and to suppress activity from about 2-4 hours after the start of inhalation.

**Supplementary Analysis of the Sex Differences in Adolescent Temperature:**

The main study was not powered to evaluate secondary factors such as the rate of return to normal temperature across sex. As an exploratory analysis the temperature responses of the two sexes on Day 1 vs Day 4 of the first week were compared. These days are the ones identified in our Nguyen et al 2018 report, using rectal sampling on adult rats, to index a sex difference in tolerance. A three-way ANOVA with repeated factors of Day and Time identified a significant effect of Day [F (1, 70) = 165.0; P<0.0001] and of Time [F (9, 70) = 13.91; P<0.0001], as well as the interactions of Time with Day [F (9, 70) = 4.33; P<0.0005] and of Sex with Day [F (1, 70) = 68.72; P<0.0001]. Evaluation of the post-hoc comparisons confirmed that for the males, temperature was not different between the Day 1 and Day 4 at any time after vapor initiation. For the females, temperature was different between the two days from 60-240 minutes. The post-hoc comparison of the cells relevant to the question of differential tolerance did not confirm any significant difference between male and female body temperature on Day 1 for any timepoint. It did confirm that male and female temperature differed on Day 4 from 60-90 minutes after the start of inhalation, the same points where the males differed from their own PG comparison day and the females did not, as per the analysis in the main results.

The second exploratory analysis contrasted the temperature response of both sexes on PG and all five THC inhalation days of the first [Time, F (1.970, 165.5) = 138.8; P<0.0001; Sex/Day, F (11, 84) = 10.71; P<0.0001; Interaction, F (99, 756) = 6.52; P<0.0001] and second [Time, F (2.271, 190.7) = 79.15; P<0.0001; Sex/Day, F (11, 84) = 5.63; P<0.0001; Interaction, F (99, 756) = 4.36; P<0.0001] weeks. The post-hoc analysis confirmed that the marginal means (i.e., the day summary) for Days 1-5 do not differ from each other in the male group. In the female group, Day 1 differs from all the subsequent days, Day 2 and Day 3 differ from Days 4-5 but not each other and Day 4 and 5 do not differ. The average temperature differs between the sexes in the PG baseline and on Day 1, Day 4 and Day 5, but not on Day 2 or Day 3. In the second week, Days 9-10 do not differ from each other in the male group. In the female group, Day 7 differs from Days 8 and 10 but the rest of the comparisons between days do not differ. Across sex, Days 6, 7 do not differ, however there is a sex difference on Day 8, Day 9 and Day 10.

**Effect of repeated THC or PG inhalation on bodyweight in adolescent rats:**

The body weight of the two male groups differed during chronic vaping (**Figure S3**). The AVOVA confirmed significant effects of Group [F(1,14)=7.95; *p*<0.05], of PND [F(19,266)=2001; *p*<0.0001] and of the interaction of factors [F(19,266)=9.35; *p*<0.0001]. The post hoc test further confirmed significant differences between groups on PND 42-45, PND 51 and PND 65. The body weight of the two female groups did not differ during chronic vaping. Although the ANOVA confirmed a significant effect of PND [F(19,266)=972.5; *p*<0.0001] and of the interaction of Group with PND [F(19,266)=2.672; *p*<0.0005], the post hoc test did not confirm any significant differences between groups for any Day. There was, however, a difference in the weight gain trajectory across chronic treatment weeks. The THC group weighed significantly more on PND 36-39, and the PG group significantly more on PND 37-39, compared with PND 36. In contrast, only the PG group gained significant weight on PND 45-46 compared with PND 42.

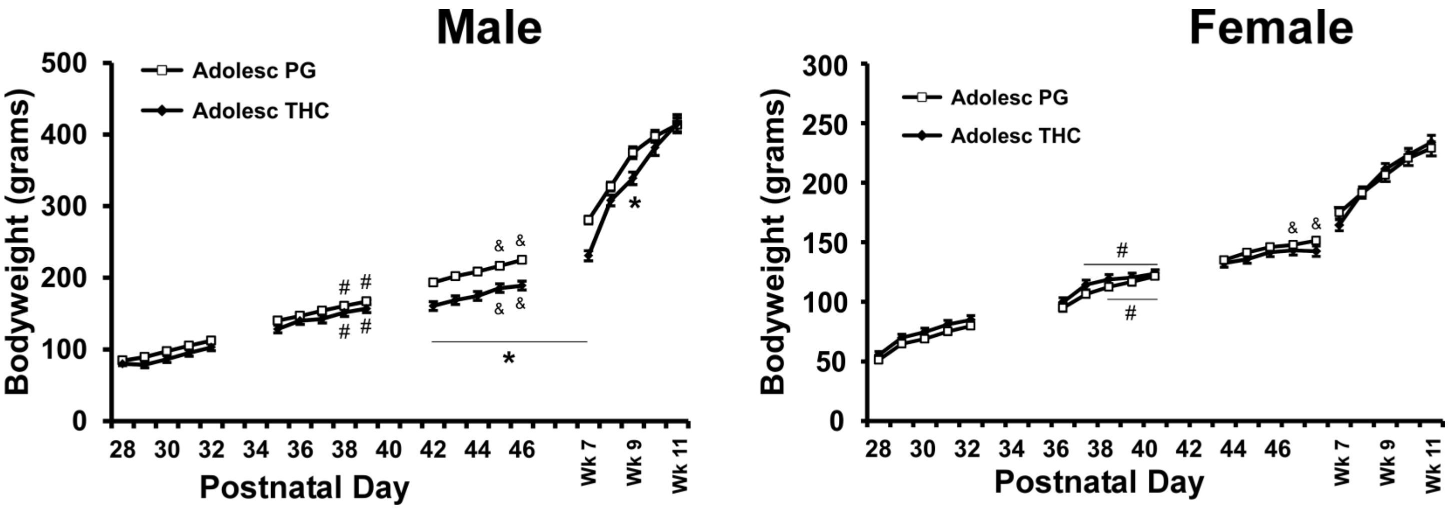


***Figure S3:*** *Mean (N=8 per sex; ±SEM) bodyweight of the repeated-PG and repeated-THC groups of male and female rats are depicted. A significant difference between groups is indicated by *. A significant increase in weight within-group relative to PND36 is indicated with # and an increase relative to PND42 with &.*

**Effect of repeated adolescent THC or PG inhalation on THC-induced nociception in female rats in adulthood:**

The female groups each exhibited slowed tail-withdrawal latency after THC inhalation, however the magnitude was reduced in the adolescent-THC group (**Figure S4**). The two way ANOVA confirmed significant effects of Group/Vape condition [F (3, 26) = 45.85; *p*<0.0001], of Time after vapor initiation [F (3, 78) = 26.04; *p*<0.0001] and of the interaction of factors [F (9, 78) = 5.23; *p*<0.0001]. The post hoc test confirmed significant increases in latency after THC inhalation for each of the repeated-PG (35-120 minutes after vapor initiation) and repeated-THC (35-120 minutes after vapor initiation) groups. The post-hoc test further confirmed significant differences between groups after THC inhalation (collapsed across time points and also 120 minutes after vapor initiation).

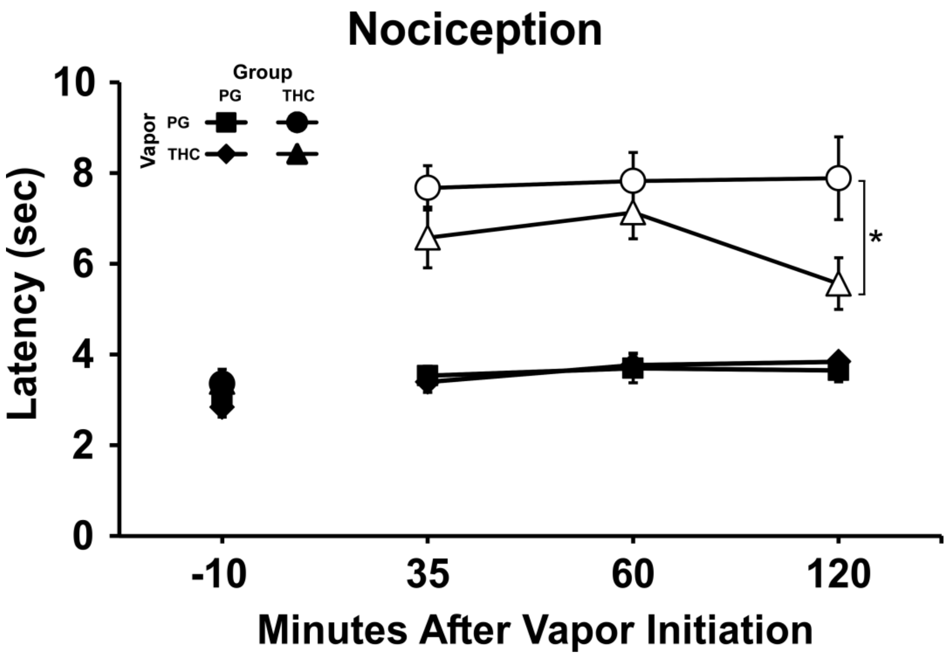


***Figure S4:*** *Mean (N=8 PG, N=7 THC; ±SEM) tail withdrawal latency following vapor inhalation of PG or THC (100 mg/mL). Open symbols indicate a significant difference from the pre-inhalation baseline and the respective timepoint after PG inhalation, within-group. A significant difference between groups is indicated with *.*

**Effect of Adolescent Treatment, Across Sex, on Oxycodone Self-Administration:**

As shown in the main text, the adolescent treatments did not alter oxycodone self-administration during acquisition or during either of the FR or PR dose-substitution experiments when considered within sex. The primary analysis did not directly compare adolescent treatment when collapsed across male and female groups, thus the following presents analysis of the effect of adolescent treatment for all rats.

**Acquisition:** The statistical analysis of acquisition by adolescent treatment, i.e. collapsed across sex groups (**Figure S5A,B**), did not confirm any treatment-group differences in oxycodone acquisition [Sessions: F (15, 555) = 25.35; P<0.0001; Adolescent Treatment: F (1, 37) = 2.06; P=0.1592; Interaction: F (15, 555) = 0.32; P=0.9935], however there was a Treatment difference in the lever discrimination [Sessions: F (15, 555) = 14.73; P<0.0001; Adolescent Treatment: F (1, 37) = 5.64; P<0.05; Interaction: F (15, 555) = 1.04; P=0.4102]; the post hoc test failed to confirm Treatment differences for any specific Session. Lever discrimination was

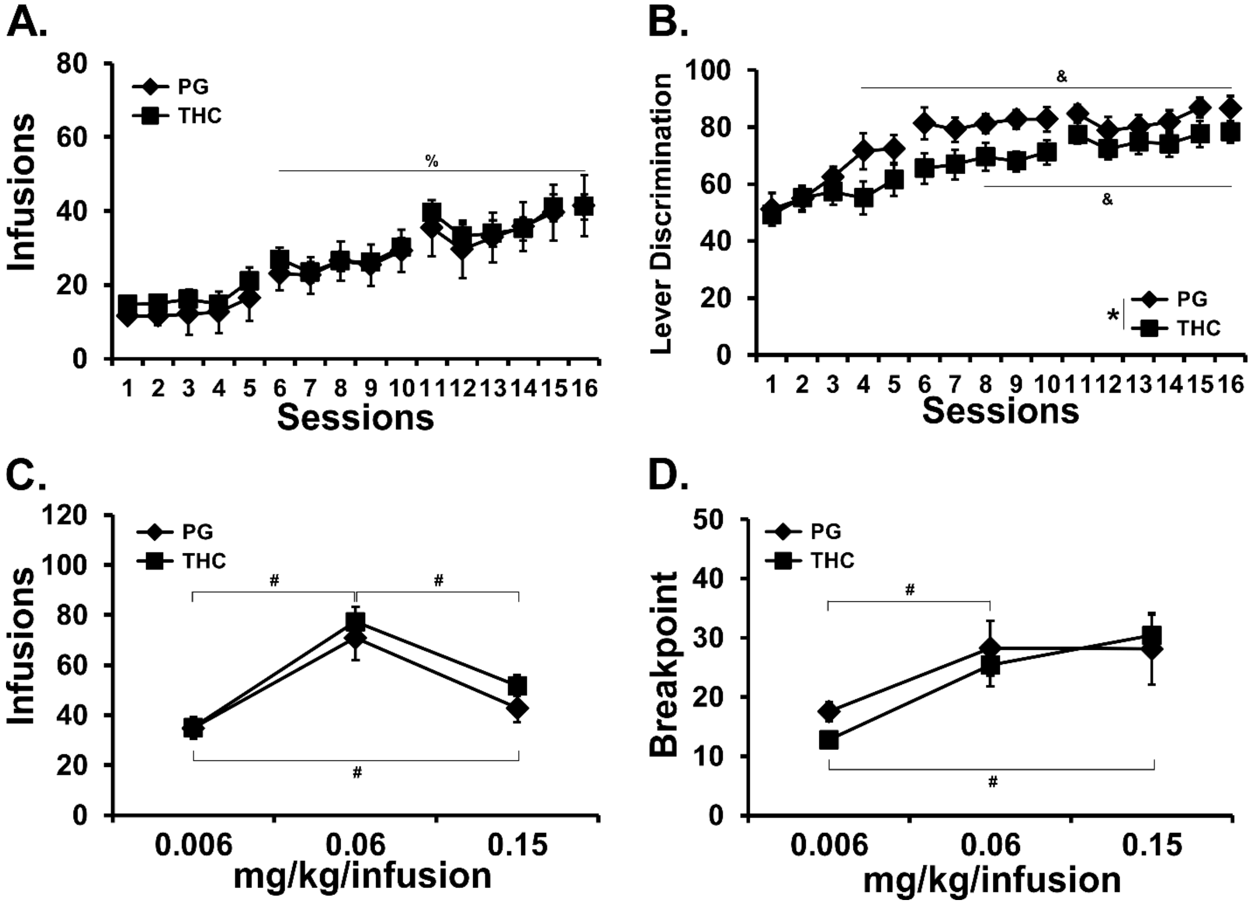


***Figure S5**: Mean (±SEM) infusions and lever discrimination during acquisition for the PG (N= 20) vs THC (N= 19) treatment groups, collapsed across female (N= 15) and male (N= 24) sex. A significant difference from the first session across group is indicated with %, a difference from the first session within group by & and a significant difference between doses, collapsed across group is indicated with #.*

affected by adolescent treatment [Adolescent Treatment: $F(1, 37) = 5.64$; $P<0.05$; Sessions: $F(15, 555) = 14.73$; $P<0.0001$; Interaction: $F(15, 555) = 1.04$; $P=0.4102$], however the post-hoc did not confirm treatment group differences for any specific session. The post-hoc test did confirm that the interaction was attributable to the THC-exposed animals exhibiting a significant change from the first session much later than the PG-exposed group.

**Dose Substitution under FR and PR:** This secondary analysis did not confirm any adolescent treatment-related effects in the FR dose-substitution [Adolescent Treatment: $F(1, 36) = 0.53$; $P=0.4735$; Dose: $F(2, 72) = 33.86$; $P<0.0001$; Interaction: $F(2, 72) = 0.44$; $P=0.6438$] and the post-hoc test of the marginal means for Dose confirmed significant differences in the infusions obtained in all three dose conditions (**Figure S5C**). In the PR dose-substitution, there were likewise no significant effects of treatment [Adolescent Treatment: $F(1, 36) = 0.18$; $P=0.6764$; Dose: $F(2, 72) = 18.74$; $P<0.0001$; Interaction: $F(2, 72) = 1.09$; $P=0.3410$] group confirmed (**Figure S5D**). The post-hoc test of the marginal mean for Dose confirmed significant differences in the infusions obtained in the 0.06 and 0.15 mg/kg/infusion condition compared with the 0.006 mg/kg/infusion dose condition.

**Female Group Treatment Between Oxycodone and Fentanyl Dose Substitution Experiments:**

For the oxycodone dose-substitution under PR (**Figure S6**), the statistical analysis did not confirm any effect of group or of oxycodone dose, there was a significant effect of pre-treatment THC dose [F (3, 66) = 10.43; P<0.0001]. There was no significant effect of adolescent treatment in the analysis collapsed across oxycodone per-infusion dose (**Figure S6A**).When collapsing the data across adolescent treatment group (**Figure S6B**) there was no significant effect of per-infusion dose but a significant effect of THC pre-treatment dose [F (3, 72) = 10.40; P<0.0001]. The post-hoc test confirmed that fewer infusions were obtained after the

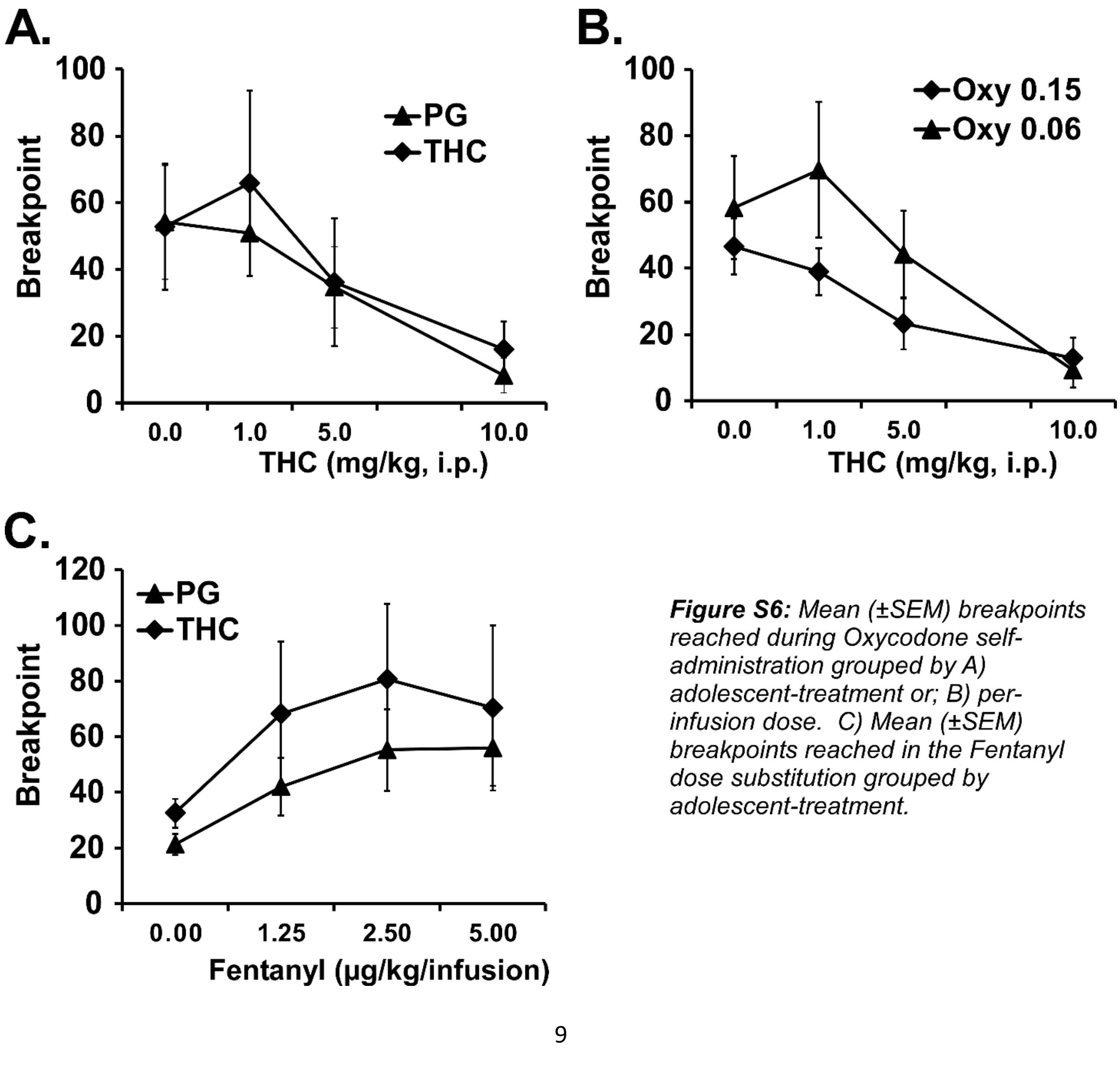


***Figure S6:** Mean (±SEM) breakpoints reached during Oxycodone self-administration grouped by A) adolescent-treatment or; B) per-infusion dose. C) Mean (±SEM) breakpoints reached in the Fentanyl dose substitution grouped by adolescent-treatment.*

10.0 mg/kg THC dose compared with vehicle and 1.0 mg/kg THC pre-treatment conditions. Next, the rats completed a PR dose-response study in which the self-administered drug was fentanyl (0.0, 1.25, 2.5, 5.0 μg/kg/infusion). There was again no significant effect of adolescent treatment group (**Figure S6C**). There was a significant effect of per-infusion dose across group [$F(3, 36) = 6.53$; $P<0.005$], but post-hoc test confirmed only that a lower breakpoint was reached in the saline condition compared with all active dose conditions.
